# Modelling single-stress omics integration with HIVE enables the identification of responding signatures to multifactorial stress combinations in plants

**DOI:** 10.1101/2024.03.04.583290

**Authors:** Giulia Calia, Sophia Marguerit, Ana Paula Zotta Mota, Manon Vidal, Hannes Schuler, Ana Cristina Miranda Brasileiro, Patricia Messenberg Guimaraes, Silvia Bottini

**Author notes:** Corresponding Author: To whom correspondence should be addressed: Silvia Bottini, PhD Institut Sophia Agrobiotech UMR 1355 INRAE/7254 CNRS/UniCA 400 route des Chappes – BP 167 06903 Sophia Antipolis Cedex France.

## Abstract

All organisms are subjected to multiple stresses usually occurring at the same time, requiring the activation of the appropriate signalling pathways to respond to all or by prioritizing the response to one stress factor. Plants, as sessile organisms, are particularly impacted by the constantly changing environment that is often unfavourable or even hostile. Because of the experimental complexity of studying the response of one organism to multiple stressors simultaneously, usually experiments are conducted considering one individual stress factor at the time. An alternative consists in performing *in silico* integration of those data on single stress response. Currently used methods to integrate unpaired experiments consist of performing meta-analysis or finding differentially expressed genes for each condition separately and then selecting the commonly regulated ones. Although these approaches allowed to find valuable results, they mainly identify specific signatures in response to one stress and very few signature responding to multiple stresses and lack those modulated differently in each condition.

For this purpose, we developed HIVE (Horizontal Integration analysis using Variational AutoEncoders) to integrate multiple single-stress transcriptomics datasets composed of unpaired experiments. Briefly, we coupled a variational autoencoder, that alleviates batch effects, with a random forest regression and the SHAP explainer to select relevant genes modulated specifically in response to one or multiple stresses.

We illustrate the functionality of HIVE to study the transcriptional changes of several different plants namely *Arabidopsis thaliana*, rice, maize, wheat, grapevine and peanut by collecting publicly available experiments on single stress, either biotic and/or abiotic, and jointly analyse them. HIVE performed better than the differential expression analysis, meta-analysis and the state-of-the-art tool for horizontal integration allowing to identify novel promising candidates responsible for triggering effective defence responses to multiple stresses.

## Introduction

From omics to multi-omics, we have been assisting to a wide growing of data production with a multitude of experimental designs. These studies can provide a global understanding of the flow of information, by allowing to infer the complete network of interactions from the original biological process to the functional consequences (1). In the last decade, omics and multi-omics studies have been employed in several biological fields, including the study of plant immunity (2, 3). Plants exhibit remarkable cellular plasticity during their growth and development, for instance, they are able to repair tissues after wounding or to regenerate entirely new organisms, as well as to reprogram cells to aid symbiotic interactions with other organisms or to counteract the attack of bio-aggressors (4). Similar to many biological processes, the plant defence response mechanisms are composed of several physiological and molecular changes mediated by complex regulatory interactions. Those processes occur at different biological layers (such as post-transcriptional, post-translational, translational and transcriptional changes), depending on cells or tissues and the several spatio-temporal scales. During the infection process, pathogen proteins play a crucial role in the re-wiring of multiple biochemical processes occurring in the host, ultimately allowing for the infection’s progression. Parasites modulate the immune response, cell signalling, MAPK and hormone signalling, vesicle trafficking systems, ubiquitination systems, metabolism or even plant growth (5). As a counterpart, hosts protein machinery trigger defence mechanisms against the pathogen. Furthermore, as sessile organisms, plants must endure abiotic stresses such as drought, high or low temperature, nutrient deficiency, salinity that can occur concomitantly to pathogens attacks. All those conditions affect plant physiology causing several changes in molecular processes that can be non-adaptative or adaptative responses (6). The former are direct responses to the stress, for instance protein structure changes caused by extreme temperatures or disruptions in enzyme kinetics and molecular interactions caused by toxic ions. The second category concerns responses that can improve stress resistance including the repair of stress-induced damage, the rebalancing of cellular homeostasis and the adjustment of growth (7, 8). Those represent potential targets for crop improvement. Therefore, understanding how plants perceive stress signals and adapt to adverse environmental conditions, both biotic and abiotic, is critical for global food security.

Although in real life multiple stresses occur contemporarily, omics experiments mimicking those conditions are hard to perform. Hence usually, omics experiments are performed by studying the organism’s response to one stress at the time and then looking for common or specific responding signatures after the data analysis. The limitation of this approach consists in an underestimation of inter-conditions information yielding a very low number of molecular signatures modulated in response to multiple conditions. Hence, there is an exponential trend in terms of volumes and complexity of omics data that are currently under-exploited due to the absence of a methodology to successfully integrate data from different experiments. Multiple challenges need to be addressed (9, 10). On the top of all, the notorious batch effects caused by external factors may hinder a joint analysis (11, 12). Batch effects are common technical variations in omics and multi-omics data and lead to equivocal results if uncorrected or over-corrected (13–16).

Lately, many techniques have been developed to address the need for batch effect correction; however, they have been developed mainly for large-scale settings (14, 17–21). In a recent study, Yu et al. (2023) showed that the best-performing batch correction method is the ratio-based technique. The drawback of this method is the need for reference samples not always available. Several studies applied autoencoder (AE) or its variants to remove the batch effects in single-cell omics and multi-omics data achieving good performances (22–24). The advantage of AEs is their powerfulness to capture non-linear patterns from high-throughput data by transforming them in a lower dimension called latent space. This new compressed representation of the original data has the limitation, prevalent among deep-learning models, of the lack of explainability because the contribution of the originally measured molecules to the latent features is not straightforward. Although some tools have been proposed, such as DeepLIFT (25) and DeepSHAP (26) they are usually very computationally intense. Furthermore, they have mainly been used coupled to a supervised approach in which the latent space is used as input for a classifier model to classify samples (i.e. cancer classification). To circumvent the needs for batch effect correction, the most commonly used approach is a meta-analysis of each experiment separately and the subsequent selection of the most common differentially expressed genes. Among such methodologies, Miodin (27) and metaRNAseq (28, 29) are two distinct packages in R language that implement statistical solutions to perform a multi-differential expression analysis and associate a p-value that considers the multiple testing. The major pitfall of this analysis is the small number of common signatures, which decreases with increasing number of datasets. An alternative is to use integrative approaches that simultaneously analyse all the experiments. In this regard MINT is an approach to analyse unpaired experiments based on partial-least square discriminant analysis (30) that corrects for batch effects, classifies samples and identifies key discriminant genes (31). However, the employability of this tool is limited to the integrations of multiple experiments all studying the same conditions thus preventing to integrate multi conditions from different experiments. Furthermore, the use of the supervised classification of samples when the experimental design includes the presence of many classes, and few replicates could affect the performance of the tool.

To address these limitations, we developed HIVE (Horizontal Integration analysis using Variational AutoEncoders) to analyse at the same time unpaired multi-conditions transcriptomics experiments. HIVE is particularly suitable when multiple conditions, each from a different experiment, need to be analysed jointly to capture common and/or specific molecular signatures. This peculiar scenario is very common in plant biology because multi-stress experiments are complicated to perform. Therefore, HIVE will provide the possibility to perform multi-stress joint analysis *in silico*. We show the capabilities of HIVE on several datasets composed each of different biological conditions and transcriptomic experiments.

## Materials and Methods

### Datasets collection, preprocessing and integration

We collected RNA-sequencing data from Expression Atlas database and previous publications (32), and microarray data from PlaD database (33) for a total of seven plant species and 77 experiments. Specifically, form Expression Atlas we retrieved three experiments from *Arabidopsis thaliana* plants accounting for five conditions and three experiments from *Vitis vinifera* plants (cv. Chardonnay and Tocai) for a total of eight conditions. In both cases RNA-seq data were retrieved from healthy and affected tissues under the attack of different biotic stressors, either environmental ones, like insect feeding, or infections. We also extracted five experiments from *Zea mays* plants representing a total of eight conditions, namely biotic stresses (environmental or infection related), abiotic stresses (wounding) and a combination of the two stresses. Finally, we also collected six different experiments from *Arachis spp. (A. duranensis* and *A. stenosperma*), retrieved from previous publications (34–37). The available conditions are: biotic (root knot nematode infection), abiotic (drought) stress, and traumatic abiotic stress (desiccation) for both plants species, and the cross-stress (biotic and abiotic at the same time) only for *A. stenosperma* plants, for a total of seven conditions. In further analysis, apart from the construction and characterization of the VAE, only nematode infection, drought and cross-stress will be taken into consideration as biotic, abiotic and cross-stresses, for a total of five conditions (refer to Table 1 and Supplementary Table 1 for detailed information on conditions). From PlaD we obtained 30 experiments from *Arabidopsis thaliana*, eight from *Oryza sativa*, 15 from *Triticum aestivum* and seven from *Zea mays* plants, under biotic stresses although no detail about the specificities of the stressor and the correspondence between samples and conditions was not provided (Table1 and Supplementary Table 1).

**Table 1.** Integrated datasets information. For each integrated dataset are reported: the original source of the data, the organism on which the experiments are conducted, the number of experiments considered (batches), the number of samples retrieved from each integrated dataset and the number of considered conditions when available.

| Data origin | Plant | N. experiments | N. samples | N. conditions |
| --- | --- | --- | --- | --- |
| Expression Atlas(32) | *Arabidopsis thaliana* | 3 | 42 | 5 |
|  | *Vitis vinifera* | 3 | 77 | 8 |
|  | *Zea mays* | 5 | 101 | 8 |
| Previous publications | *Arachis spp.* | 6 | 35 | 7 (5) |
| PlaD(33) | *Zea mays* | 7 | 92 | NA |
|  | *Oryza sativa* | 8 | 185 | NA |
|  | *Triticum aestivum* | 15 | 432 | NA |
|  | *Arabidopsis thaliana* | 30 | 762 | NA |

The gene expression count matrices were available either from the used databases or from the original publications (for the procedure refer to the experiments ID or the original publications listed in Supplementary Table 1). The integration of all datasets corresponding to each plant model was performed via HIVE code, concatenating samples and replicates of each experiment, for a total of eight integrated datasets.

With a HIVE built-in function, we filtered out genes having a variance value belonging to the first quartile of the distribution. The final integrated datasets contain 24,624 genes for *A. thaliana* (RNA-sequencing experiments), 22,478 genes for *V. vinifera*, 32,227 genes for *Z. mays* (RNA-sequencing experiments), 27,557 genes for *Arachis spp*, 15,423 genes for *A. thaliana* (microarray experiments), 15,082 genes for *O. sativa*, 45,964 genes for *T. aestivum* and 9,956 genes for *Z. mays* (microarray experiments).

### HIVE framework overview

HIVE workflow consists of three main blocks described in the next paragraphs and summarized in Figure 1. Briefly, the first block concerns horizontal integration of datasets and batch effect correction. A Variational AutoEncoder (VAE) is used to encode the integrated dataset in a compressed representation called latent space. The two following blocks are independent from each other, but they are consequent to block 1. One block regards the exploration of the latent features. Therefore, the point biserial correlation and the Kruskal-Wallis test followed by the post-hoc Dunn’s test are applied to calculate the association between the latent features and phenotypic characteristics(38–40). The last block is composed of four steps to select relevant genes for the studied phenotype. First, a Random Forest Regression (RFR), in a k-fold cross-validation configuration is used where each latent feature is considered as the variable to explain, and the original gene expression data as the dependent variable. The problem of inferring the relationship between a set of N latent features and G genes is decomposed into a set of N regression problems. Afterward, the contribution of each gene to each regression was calculated with the SHAP algorithm. Finally, we included a custom method to set a threshold on the SHAP scores to select only the most contributing genes. The last step consists in associating the genes to the most varying condition(s) compared to each control. At the end, among other meta-outputs, HIVE returns a list of ranked genes that can be used for further analysis.

**Figure 1.**
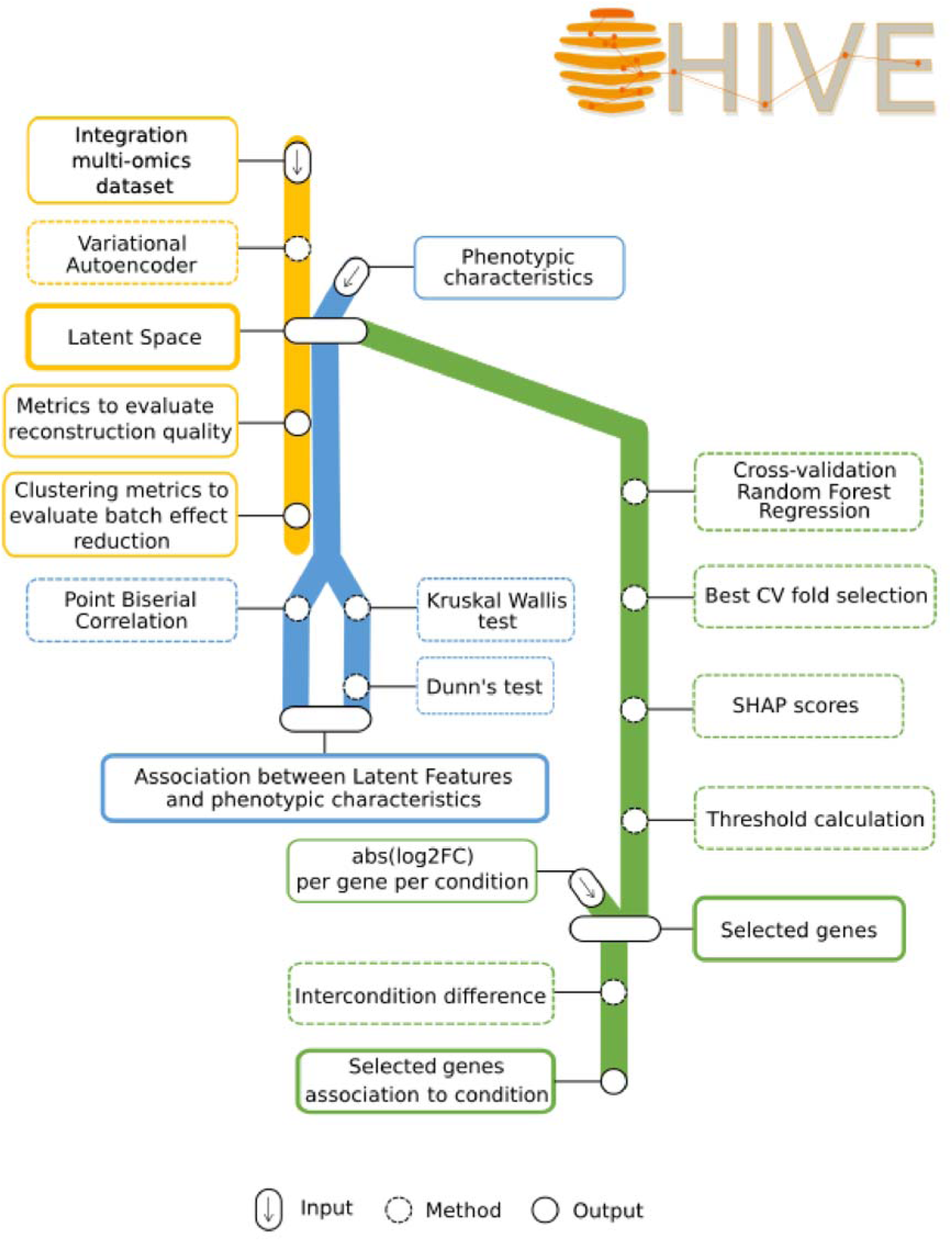
HIVE framework. The pipeline is represented as three underground lines to symbolize the three main blocks composing HIVE. The first block (yellow line) concerns the batch effect correction using a Variational AutoEncoder (VAE) to produce the latent space, that is the common “station” of all the lines. The two following lines are independent of each other, but they are consequent to block 1 because they use the latent space. The blue line regards the calculation of the association between the latent features and phenotypic characteristics using the point biserial correlation. The green line is composed of four steps to select relevant genes for the studied phenotype. First, a Random Forest Regression (RFR) is used where each latent feature is considered as the variable to explain, and the original gene expression data as the dependent variable, in a cross-validation framework. Afterward, the contribution of each gene to each regression was calculated with the SHAP algorithm. We included a custom method to set a threshold on the SHAP scores to select only the most contributing genes. The third step returns a list of ranked genes that can be used for further analysis. Lastly, by implementing a novel method based on the amplitude of difference in expression value, each gene is associated to one or more conditions.

### Block 1: batch effect correction

#### Variational Autoencoder configuration

A VAE is used to encode the integrated dataset in a compressed representation called latent space. An Autoencoder is a type of deep neural network which is trained in a self-supervised manner (41, 42). The symmetric structure is composed of an encoder, a decoder and a bottleneck in the middle which we will refer to as latent space. The network is therefore trained to reconstruct the input data with minimal error by learning to encode the N-dimensional data as a L-dimensional latent space, with L<<N. The VAE originally proposed by Kingma and Welling (43), and Rezende et al. (44), can be seen as a probabilistic version of an AE, where the resulting model can be also used to generate new data. The encoder is trained to estimate the mean and the variance model by a multivariate gaussian mixture. Briefly, the latent variables Z are assumed to have distribution *pθ* and the observation X is assumed to be conditional distribution *pθ(x|z).* The generative model is trained to estimate the parameters *θ* to minimize the Kullback-Leiber (KL) divergence between the original data distribution and the model distribution (the marginal likelihood). Being the marginal likelihood analytically intractable because of the nonlinear relationship between the original variables and the latent components, the evidence lower bound (ELBO) approximation is used. The VAE architecture, performances and robustness are fully described in the Supplementary Note section of Supplementary Material and Supplementary Note Figure 1, 2 and Supplementary Note Table 1, 2. A min-max transformation was applied on raw transcriptomics data before applying the VAE, using MinMaxScaler function of Scikit-learn python library (45). The entire VAE is written in python language and uses Tensorflow and Keras libraries (46, 47). The output of this first step is the latent space as a matrix having 80 latent features as columns and samples as rows.

**Figure 2.**
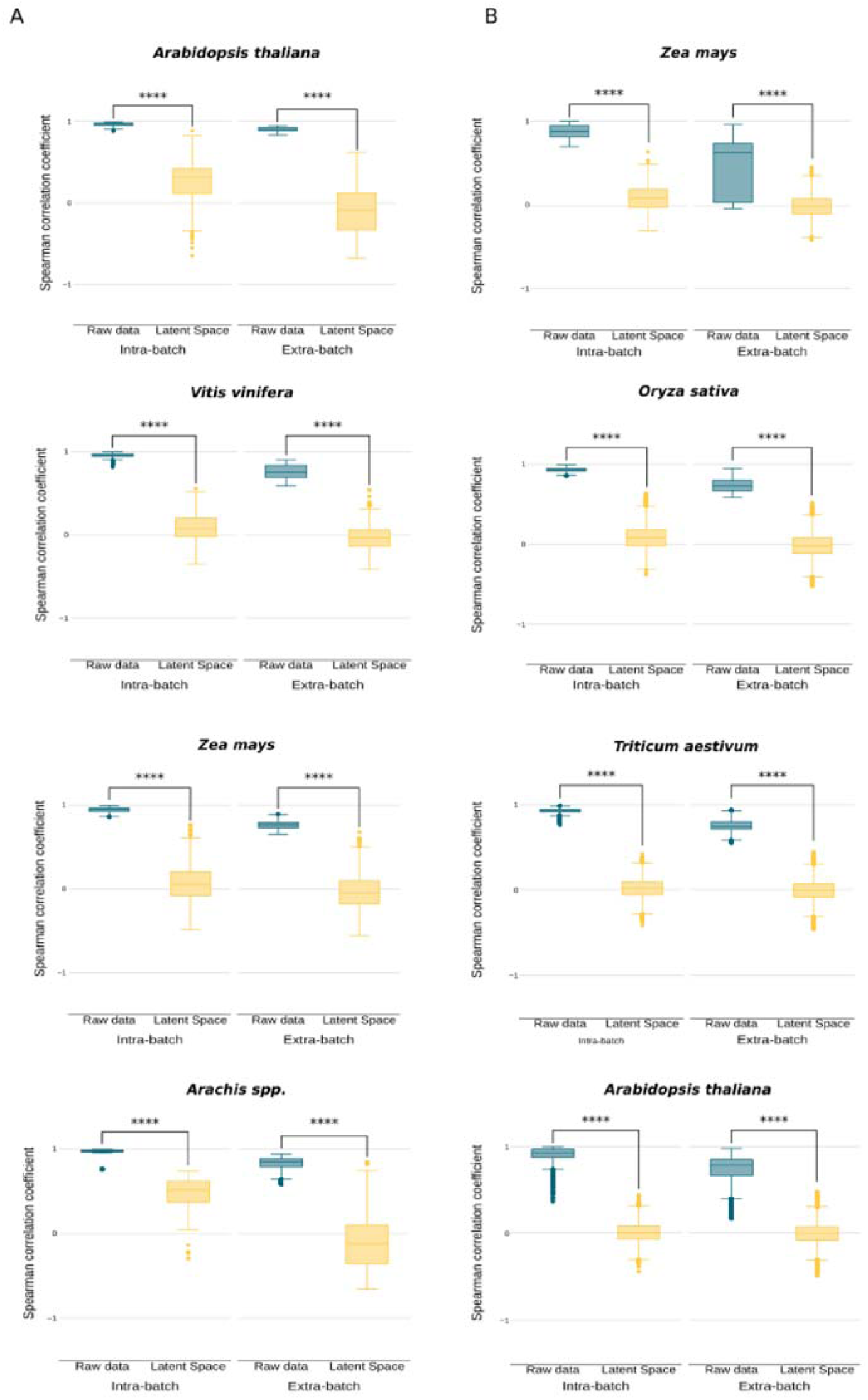
Intra- and Extra-batch correlation of samples. A) Spearman correlation between samples in datasets from RNA sequencing data. The “****” represents a significant p-value < 1.3e-19 for Wilcoxon test. B) Spearman correlation between samples in datasets from microarray data. The “****” represents a significant p-value < 1.3e-160 for Wilcoxon test.

### Block 2: Association between latent features and phenotypic characteristics

To calculate whether the latent space was able to capture phenotypic characteristics, we implemented two methods. The point biserial correlation (p-value <= 0.01), a correlation coefficient measure, that can be applied when one of the variables is dichotomous (when the phenotypic variable have only two characteristics) and the other is continuous (the latent features) (38). Assuming to have two sets of elements to correlate, i.e. X = {x_1_, …, x_n_} (dichotomous variable) and Y = {y_1_, …, y_n_} (continuous variable) with n the total number of observations, n_0_ the number of elements in X which are 0 and n_1_ the number of elements in X which are 1, the point biserial correlation can be calculated as follows:

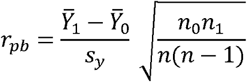

Where *Ȳ*_0_ and *Ȳ*_1_ are the means of the corresponding Y elements in X, coded 0 and 1 respectively, and *s_y_* in the standard deviation of Y. In case of more than two characteristics, we combine the use of the Kruskal-Wallis test to identify the latent features that are significantly associated with multiple-calls phenotypic characteristics, followed by the post-hoc Dunn’s test to identify specifically to which characteristic each latent feature is associated with.

We implemented this calculation in python language and using both SciPy and Scikit-learn libraries (48, 45) included in the HIVE pipeline.

### Block 3: Gene selection

#### Random Forest Regression

Before performing the Random Forest Regression (RFR), genes having a raw expression value for each sample lower than 30 are filtered out and latent features are scaled using MinMaxScaler function of Scikit-learn python library. We implemented a cross-validation strategy to train and test the regression models and identify the best performing one. Cross-validation is useful in case of limited amount of data, because splitting into training and test may yield a small test set (49–52). In case of cross validation, we can build different models to be sure to make prediction on all the data. Here we used a stratified k-fold cross-validation that allow the model to make predictions on class-proportioned unseen data for each fold. Thus, we can use the entire dataset for training and test. The number of folds depends on the minimal number of replicates per condition to assure to have balanced representation of samples in each class during this phase. For the application on our integrated dataset, we performed a three-fold cross-validation. The RFR (53) is implemented using RandomForestRegressor() method in sklearn.ensemble module from Scikit-learn python library, using default parameters.

#### SHAP scores

For each latent feature, the best performing model from cross-validation is used to calculate the contribution of each gene to each regression with the SHAP (SHapley Additive exPlanations) explainer. SHAP is a method based on the Shapley interaction index from game theory and used to increase transparency and interpretability of machine learning models (26). Briefly, the regression model can be seen as a cooperative game and the SHAP explainer assigns a score to each feature (i.e. gene) representing its contribution to the model’s output. We used the SHAP TreeExplainer from SHAP python library (54).

#### Threshold calculation and gene selection

We introduce a custom method to calculate a threshold on the SHAP scores to select only the most contributing genes to each regression (extended description in Supplementary Note). For each latent feature, we calculated the distribution of SHAP values by grouping the values in 60 ranges, each range is called bin hereafter (Supplementary Note Figure 3). The number of bins is a modifiable parameter in HIVE allowing the user to adjust this value. By setting a threshold on the SHAP values in the bins, we retrieved only the genes with SHAP scores in the tails of the distribution, therefore those with the most extreme SHAP values and indicating a major contribution to the corresponding latent feature.

**Figure 3.**
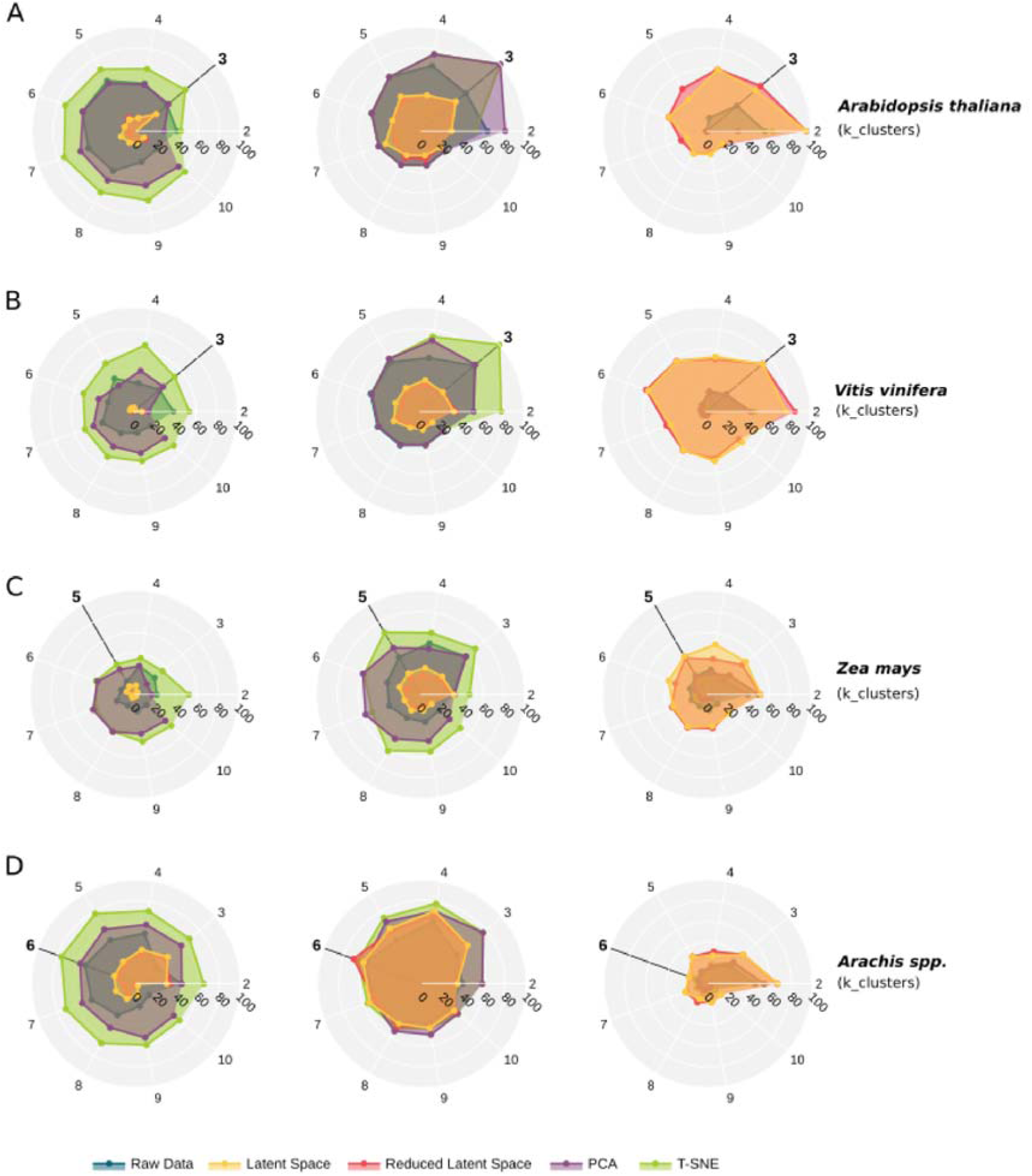
Clustering performances for the benchmarked RNA sequencing. Radar plots summarizing the agreement between of K-means clustering with increased number of clusters (*k*) and batch belongings with three metrics, Silhouette score, Jaccard index and entropy for A) *Arabidopsis thaliana* B) *Vitis vinifera* C) *Zea mays* and D) *Arachis spp* datasets respectively. The number of original batches in the considered dataset is highlighted in the correspondent plot.

After repeating the procedure for each latent feature, HIVE provide a unique list of ordered genes accordingly to their SHAP value rank. If a gene is selected by more than one feature, then only the highest rank will be maintained in the final list.

#### Gene association to condition

To identify in which condition(s) the HIVE selected genes have the highest deviation from the respective control sample, we set up a method based on the amplitude of variation. First, we calculated the logarithm to base two of the fold-change (log2FC) for each condition with respect to the control using the rlog function in the DESeq2 package (R v4.1.2) (55, 56).Then for each gene we use the highest absolute value as reference and calculated the distances with each of the other conditions. By selecting as threshold half of the standard deviation of the total distance distribution (Supplementary Note Figure 3D). We consider each gene to be associated to the condition with the highest abs(log2FC) and to all other condition(s) that have a distance smaller than the selected threshold.

### Metrics for benchmarking batch effect correction

To evaluate the batch effect correction ability of the HIVE VAE, we first compared the batch effect in both raw data and obtained latent space for each of the eight integrated datasets. With this purpose we calculated the Spearman correlation coefficient of samples in the integrated raw matrix and in the latent space, using the R “cor” function, and then compared the intra-batch effect, namely the distribution of samples’ correlation coefficients belonging to the same batch and the extra-batch effect, namely distribution of samples’ correlation coefficients in one batch compared to all the other batches. To verify the difference between the coefficient distributions we used the Wilcoxon signed-ranked test (p-value<= 0.05), from python library SciPy.

Afterwards we compare the HIVE-VAE with other commonly used dimensionality reduction techniques namely Principal Component Analysis (PCA) and t-distributed Stochastic Neighbour Embedding (t-SNE). We standardized the data using the StandardScaler() from Scikit-learn python library. For the PCA we considered 10 principal components representative of most of the variance in the data.

As first analysis, for each configuration (raw-data, VAE, PCA, t-SNE), we performed a K-means clustering with k varying accordingly to the number of batches in the integrated dataset. We measured clustering performances in terms of: homogeneity score, completeness score, v-measure, adjusted Rand score, adjusted mutual score, Jaccard score, silhouette score (Bray-Curtis metric), inertia and entropy. For most of them, we used the module ‘metrics’ of the python library Scikit-learn, but we implemented in-house the Jaccard score and the entropy percentage, to match the need of comparing real batch labels with clustering labels. The clustering performances are included as meta-output table in every HIVE application.

As second analysis, we performed a multinomial logistic regression model implemented with python library Scikit-learn to link the batch effect to the latent components of each method. In a stratified 2-fold cross validation configuration, each variable from either the VAE latent space (latent features), or principal components from PCA or t-SNE, were used as the independent variable and a multi-class vector containing labels for batches is used as the dependent variable. The association between latent features/components and batches is evaluated by F1-measure and, for RNA-sequencing dataset we explored the mean difference of F1-score distributions via the Cohen’s effect size for HIVE and PCA application. To assess the batch effect reduction after a possible feature selection in the latent space, we then removed from the latent space only the first 8 latent features having the highest F1-measure score, representing the 10% of the latent features with higher association to batch effect, producing a “reduced” latent space.

### Metrics for benchmarking methods performances towards gene selection

We compared HIVE performance in gene selection with other two tools DESeq2 R package and the Multivariate INTegrative method (MINT) framework within the MixOmics R package (55, 57). DESeq2 is designed to calculate differential gene expression from high-throughput data, based on a negative binomial distribution. MINT is a supervised method performing horizontal integration of unpaired data and feature selection (31). For the *Arachis spp*. dataset we also compared our tool performances with the meta-analysis performed by Mota et al. (37). The meta-analysis in the original publication was performed by using the metaRNASeq R package (29).

For the three datasets from Expression Atlas database, DESeq2 results were provided for all conditions, with the exception of experiments with multi-factorial stresses in *A. thaliana* and maize plants. In those cases, we performed DEseq2 analysis with default parameters. To obtain the final list of significantly differentially expressed genes for all datasets, we retrieved all genes with p-value threshold <= 0.05 and an absolute log2FC >= to 1 in at least one condition. For MINT application, we used the sparse Partial Least Squares - Discriminant Analysis (sPLS-DA) model implemented by the tool. For the meta-analysis we retrieved the results from Mota et al, 2021 (37).

### Sharedness and specificness indexes

To evaluate the proportion of specific and common signature found by each method, we defined two novel indexes:

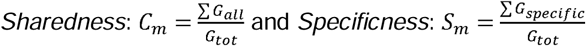

Where *G_all_* is a gene found to be modulated in all condition, *G_specific_* is a gene associate to only one condition and *G_tot_* is the total number of genes found by each method. *m* stands for method, meaning that each index is calculated for each method included in the benchmark.

### Cumulative distributions

From each list of gene found deregulated specifically to respond to one condition accordingly to each method, namely DESeq2, MINT, HIVE and meta-analysis (only for peanut dataset), we built a cumulative distribution to compare the global gene expressions of those genes with the background, i.e. all gene expressions. The python numpy.histogram() function (58) was used to first bin the distribution and count the occurrences per bin, then we calculated the Probability Density Function (PDF) and consequently the Cumulative Distribution Function (CDF) with numpy.cumsum(). To quantify the differences between the underlined distributions we used the Kolmogorov-Smirnov test (p-value <= 0.05), using the kstest module from SciPy python library.

### Functional analysis

The web-based Gene Ontology (GO) toolkit for Agricultural Community, AGRIGOv2 (59) was used to retrieve the GO-terms enrichments for gene selection of each benchmarking tools and for each RNA-sequencing datasets. In AGRIGOv2, we selected the “TAIR10_2017” gene ID types for *A. thaliana*, the “Gramene Release 50” for *V. vinifera*, the “Maize v5 (Pannzer)” for *Z. mays*, and the “Gene Model Name (PeanutBase)” for *Arachis spp.*. GO term enrichment is performed using the Hypergeometric statistical test (p-value <= 0.05), and using the Plant GO slim background ontology, developed by The Arabidopsis Information Resource (TAIR) (60).

To retrieve information such as hormone-related genes and kinases, we performed protein annotation with MERCATOR4 (61) web-based tool. First, CDS sequences were downloaded from PeanutBase database (62). The obtained sequences accurately parsed to retrieve only the HIVE selected gene sequences are then used as input of MERCATOR4. Here we specified the type of sequence as DNA and we flagged the option to include Swissprot annotations.

The list of NBS-LRR genes was retrieved from two previous publications (35, 37), including the experimentally validated. The list of transcription factors was retrieved from PlantTFDB.

All the output files have been downloaded and then parsed with *ad hoc* python scripts.

### Pathway relevance score

We defined a score for each GO term, the pathway relevance score (PR-score), calculated as follow:

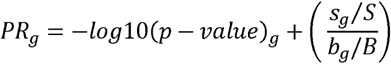

Where *g* is the considered enriched GO term, *s_g_* is the number of genes associated to the GO term *g* in the gene-selection list of interest with length (*S*), *b_g_* is the number of genes associated to the GO term *g* in the background, and *B* is the total number of coding genes in the plant of interest. Therefore, this score considers the enrichment of the related pathway and the proportion of genes associated with it on top of the original gene list.

### Coefficient of variation

The coefficient of variation is calculated on the log2FC of the genes in the considered condition/s for each enriched GO term. This is calculated by calculating the ratio between the variance of the log2FC values across all genes (calculated with “var()” method of the numpy python library) and their average log2FC.

### Voronoi diagrams

Using Voronoi diagrams we can divide the plane on which a defined number of points are laying by applying the concept of nearest-neighbouring; each point will be restricted to the region of the plane that equally separates it from all its 1-nearest-neighbours (63). To define the region, the algorithm proceeds pairwise, calculating the perpendicular bisector of each couple of points, such that the crossing of the bisectors will be the vertex of the Voronoi region forming a convex polygon (64–66). We implemented the Voronoi diagram using the classes Voronoi and Voronoi_plot_2d of the module spatial of python library SciPy. The Voronoi diagram is used, here, to represent the plethora of possible gene expressions depending on the number of conditions each gene was found deregulated accordingly to HIVE selection. Therefore, each point represents a gene, and the position depend on the level of sharedness, i.e. number of conditions in which the correspondent gene was found deregulated on the y-axis and the relative gene expression on the x-axis. For each Voronoi region including each point, we calculated the volume with the class ConvexHull of the module spatial of python library SciPy and coloured the regions accordingly. Since the points of Voronoi diagrams at the external borders of the distribution are assigned to not-finite Voronoi regions, we introduced three concentric and square-shaped mock distributions of points on the plane: the first one around the original data depending on their distribution, the second one is larger and only depends on the maximum expression of the data, and the third one is totally independent from the original data. The Voronoi regions formed by these mock points will have an assigned colour corresponding to 1 + the maximum volume of the set of original Voronoi regions, to avoid misleading-coloured regions. Finally, to compare the points density in the diagram, we defined five ranges of log2FC: from -10 to -5, from - 5 to -1, from -1 to 1, from 1 to 5 and finally from 5 to 10 and plotted the corresponding volumes distribution of Voronoi regions in the corresponding interval. We compared the distributions in each range using the Mann-Whitney U test (p-value<= 0.05), from python library SciPy.

### Hierarchical clustering of gene expression profiles

To visualize the expression profiles as heatmap and perform hierarchical clustering we used the pheatmap R package (67) using Spearman correlation as distance measure for genes clustering using as input the log2FC calculated previously.

### Gene regulatory network (GRN) inference and analysis

We used GENIE3 R package (68) to infer the GRN, providing as “regulators” the list of transcription factors and as “target” the remaining genes. We retrieved the transcription factors from *Arachis duranensis* in Plant Transcription Factor Data Base (PlantTFDBv5.0) integrated in Plant Transcriptional Regulatory Map (PlantRegMap) (69, 70). We used the Networkx python library to build-up and visualize the resulting GRN (71).

Specifically, we used the Networkx built-in function, from_pandas_edgelist(), to declare the regulatory and target genes from a pandas dataframe obtained by parsing GENIE3 output and the DiGraph() module to reconstruct a directed network. The visualization via Networkx is obtained by the built-in function draw() and specifying the node position passing to the function spring_layout() an indirect version of the previously obtained network.

We used the Leiden community detection method, implemented into the python library leidenalg that is built on the igraph library, to find communities in our network and the library on the GRN with the partition_type parameter set as leidenalg ModularityVertexPartition. We then explored each community with *ad hoc* python scripts using methods for functional analysis illustrated in the previous paragraphs.

### Overexpression of NLRs in Arabidopsis

The 1,920 bp coding sequence of *AsTIR19* from *A. stenosperma* used here was previously cloned in the pPZP_BAR *Agrobacterium* binary vector, and successfully overexpressed in transgenic tobacco plants for *Sclerotinia sclerotiorum* resistance (72).

To identify the complete coding sequence of *AsHLR*, the Aradu.HLR71 gene model from *A. duranensis* was aligned with the four best BLASTn hits of *A. stenosperma* databases available on NCBI. The consensus sequence (1,296 bp) was synthesized and cloned (Epoch Life Science Inc., TX, United States), under the control of the *Arabidopsis thaliana* actin 2 promoter (ACT-2) and the nopaline synthase (NOS) terminator, at the *XhoI* restriction site of pPZP_BAR (73).

Both vectors were used for A. thaliana ecotype Columbia (Col-0) transformation using the floral dip method (74), and the overexpression of the transgenes in the OE lines at T2 generation was confirmed by qRT-PCR analysis (75).

### Assessment of the effect of NLR genes on RKN resistance

For the *M. incognita* bioassay, four weeks old *A. thaliana* plants overexpressing *AsTIR19* and *AsHLR* genes were challenged with 500 J2 of *M. incognita* as previously described (Mota et al. 2019). Ten plants from each of the six Arabidopsis OE lines and the non-transgenic control (WT) were used, and after 60 days of infection (60 DAI), the number of eggs per gram of root (N° eggs/gr of root) was estimated in OE lines and WT plants, according to (76). The data was statistically evaluated using the Tukey test (p-value<0.05).

### Data availability

Data, scripts, and HIVE source code are available in the GitHub repository: https://github.com/Plant-Net/HIVE.git

## Results and Discussion

### Overall study design

The main objective of this work is to show that, by re-using publicly available transcriptomics data, integrative analysis with HIVE can highlight novel signatures and biological insights which cannot be found by other analysis methods. Although HIVE can be applied in any contest involving multi-transcriptomic data, here we focused to study plants respond to multiple stresses. Due to the complexity to realise in the laboratory multi-factorial stress experiments occurring at the same time or sequentially, our hypothesis is to realise multi-factorial stress experiments *in silico* by analysing omics data on single stress in an integrative fashion. By using publicly available data from Expression Atlas, PlaD or from publications, we constituted eight multi-stress datasets of transcriptomic data from either RNA-sequencing (RNA-seq) or microarray, for six different plants including: maize, rice, wheat, grapevine, peanut and *Arabidopsis thaliana* (Table 1). Importantly, for four datasets, among all the conditions, we also dispose of multifactorial stress experiments in which the plant is subjected to two stresses contemporarily (either biotic and abiotic or two different biotic agents) and the plant subjected to only one of the two at the time. Having for those biological models, both single stresses and the combined experiments, allowed to test the capability of HIVE to find signatures common to both stresses when only single-stress experiments are available.

First, we tested the performances of HIVE to reduce the batch effect and to extract biologically relevant genes by performing an extensive benchmark against other state-of-the art tools. Then we focused on the peanut dataset, since this dataset was previously studied to compare the plant response to multiple stress, to show the advantages of using HIVE to perform the analysis over the method used by Mota et al. (37).

### The variational autoencoder reduces the batch effects allowing unpaired experiments integrated analysis

In the HIVE framework, the first step consists of a variational autoencoder (VAE) based method to alleviate the batch effects and to allow the integrated analysis of unpaired experiments. VAE is a deep neural network model that learns a nonlinear mapping function from the original data space to a low-dimensional feature space called latent space (43, 77). To show the presence of batch effects in the raw data and to test the ability of the VAE to alleviate this phenomenon, we calculated pair-wise correlations among samples before and after HIVE application. For this analysis we used all the eight datasets composed by variable number of batches, samples and conditions. The number of batches ranges from 3 to 30 and the number of samples from 35 to 762, we assembled the datasets to avoid correlation between those two quantities (Supplementary Figure 1). We quantified the correlation of samples belonging to the same batch (intra-batches) and the correlation of samples from one batch with samples in all the other batches (Figure 2). The overall correlation of samples in each batch when considering the raw expression data is close to 1, while we observe a significant reduction when considering the latent space obtained by the VAE (p-values < 1.3e-19, Wilcoxon test). Using the raw expression values, the extra-batches correlations are still very high achieving similar results as the intra-batches correlations. The use of the VAE shows a strong reduction of the correlation among samples from different batches (p-values < 1.3e-160,Wilcoxon test). Those observations are consistent among all the eight datasets used, independently by the number of batches/samples and the technology used to measure expression data (RNA-seq or microarray).

This analysis shows the consistency of HIVE in alleviating the batch effects on integrated datasets composed of different number of batches from RNA-seq or microarray experiments.

### Quantitative assessment of the batch effect reduction

Batch effect reduction is a crucial step in data analysis of unpaired integrated datasets as discussed before. To quantitatively evaluate the batch effect reduction, we performed two main analyses using the eight datasets.

First, we performed clustering analysis of samples considering their transcriptomics profiles in the raw data and upon application of either PCA, t-SNE, commonly used techniques to perform dimensionality reduction on complex dataset, or the HIVE VAE, using the k-means algorithm. For each dataset, we chose a wide range of number of clusters always including the correspondent number of batches. Then we evaluated the distribution of batches in the clusters obtained from the four setups by using common metrics including Silhouette index, Jaccard score and the Shannon entropy (see methods). The Silhouette index measures how an object is more similar to elements in the same clusters than the other clusters. The Jaccard index allow us to measure how each cluster matches with each batch. Here we calculated the Jaccard index between each cluster and each batch and reported the average of the max value obtained for each cluster. The lower the value for those scores, the lower samples from the same batch cluster in the same group, therefore the better the method achieved batch effect reduction. While the opposite is expected for the entropy which measures the impurity of the studied variable or system. Strikingly, both the Silhouette score and the Jaccard index on the HIVE latent space show the lowest values and the highest entropy on all eight datasets, independently on the number of clusters in the chosen range for each dataset, compared to the correspondent scores obtained on raw data, PCA or t-SNE (Figure 3, Supplementary Figure 2, Supplementary Table 2). The only exception was reported for the Jaccard indexes for the peanut dataset, having similar values independently of the method used. Similar pattern is also found by the distributions of all Jaccard index for each cluster-batch association reported in Supplementary Figure 3BA and B, suggesting that the very low number of samples per batch in this dataset, (one batch has only one sample and three batches only three) have led to lower performances in term of Jaccard index. However other indexes like the silhouette score corroborate the batch effect reduction upon HIVE utilization for this dataset.

The second analysis we set up to test the capabilities to reduce that batch effect consists in using a logistic regression approach. While the clustering approach evaluates global batch effect reduction, the logistic regression is applied to each feature separately. We used the logistic regression to evaluate the classification performances in batches for each latent component obtained by the VAE, the first 10 principal components obtained using the PCA approach and the first two from t-SNE (Supplementary Table 3). We used the F1-score to evaluate the performances of the logistic regression for each component, expecting low F1-score for methods achieving the best batch effect reduction. Consistently with the previous results, we observe that also by considering each latent feature separately, HIVE achieved better results in batch effect reduction compared to the other two methods for all datasets. Average F1-scores for HIVE latent components spans from 12.6% for Arabidopsis dataset (microarray data), to 59% for maize (RNA-seq data). Overall, we observed that the average F1-score for microarray dataset tend to be less than the average F1-score for the RNA-seq datasets when they are analysed with HIVE and t-SNE while for PCA, the percentages remain quite comparable. Only exception is the *A. thaliana* dataset from microarray experiments where the F1-scores are the lowest among all datasets and all techniques. Importantly, the extremes of the F1-scores distribution over the 80 (HIVE), 10 (PCA) and 2 (t-SNE) components are always lower for HIVE compared with the other two techniques. Because of the higher batch effect relation of the RNA-seq data we further explore the differences in F1-score distribution for HIVE and PCA, demonstrating a batch effect reduction accordingly to the Cohen’s effect size, for HIVE application compared with the PCA (Supplementary Figure 3C). Then, we tested the clustering approach on the latent space after removing the top 10% of latent features with the highest F1-score, since those represent a low batch effect reduction; we refer to this as “reduced latent space”. As reported in Figure 3, Supplementary Figure 2 and Supplementary Table 2, the scores are similar to the ones obtained by considering all 80 latent components suggesting that feature selection in the latent space is not necessary to improve the batch effect reduction.

Overall, we quantitatively measured the batch effect reduction obtained with the VAE either considering all components, and considering each component separately. In conclusion, HIVE reduces batch effects independently of the number of batches, samples or the biotechnology (or RNA-seq) used, thus enabling the integrated analysis of unpaired experiments with complex experimental design.

### Benchmarking tools for gene selection from integrated datasets

HIVE uses a combination of random forest regression and SHAP explainer to extract the genes with expression profiles imbalanced in at least one condition. To evaluate the performances of HIVE, we compared with other tools available in the literature for gene selection. We used two methods each as representant of a category of methods to analyse transcriptomic data, namely: a classical differential expression analysis for each condition with the tool DESeq2 (78) and the MINT tool from the MixOmics R package (31, 57), which is a supervised approach to performing integrative analysis of unpaired experiments, with the constrain of requiring all conditions to be sampled in all experiments (see methods). From now on we will use only the four datasets from Expression Atlas or GEO, because for datasets from PlaD a detailed description of conditions and association between condition/sample is missing. For peanut datasets, we do not consider the desiccation condition for further analysis since only one replicate is available per peanut species, making the comparison with other tools fairly impossible. Since the peanut dataset was already used to perform integrative analysis, we also compared with the results obtained by the meta-analysis approach used in the study of Mota et al. (37) for this model.

First, we compared the total number of genes found by all methods on the four datasets. DESeq2 finds by far the largest number of genes deregulated in at least one condition for the four datasets, from 4722 (*A. thaliana* dataset) to 21.832 (maize dataset), although for the *A. thaliana* dataset the difference is of few genes (Figure 4A). On the other hand, MINT generally finds very few genes, from 8 (grapevine dataset) to 88 (peanut dataset). The number of genes found by HIVE is similar among the different datasets, from 4594 (*A. thaliana* dataset) to 8062 (maize dataset), with a slight increase corresponding to the increasing number of conditions in the different datasets. For the peanut dataset the meta-analysis originally performed in Mota et. al., 2021 (37), found a comparable number of genes as found by HIVE. By inspecting the overlap among those lists of deregulated genes, we observe that the overall agreement is very low (Figure 4B). Of note, despite DESeq2 finds a very high number of genes, very few are in common with the other methods. In average half of the genes found by HIVE were also found by DESeq2, while the agreement between MINT and the other two methods, depends on the dataset.

**Figure 4.**
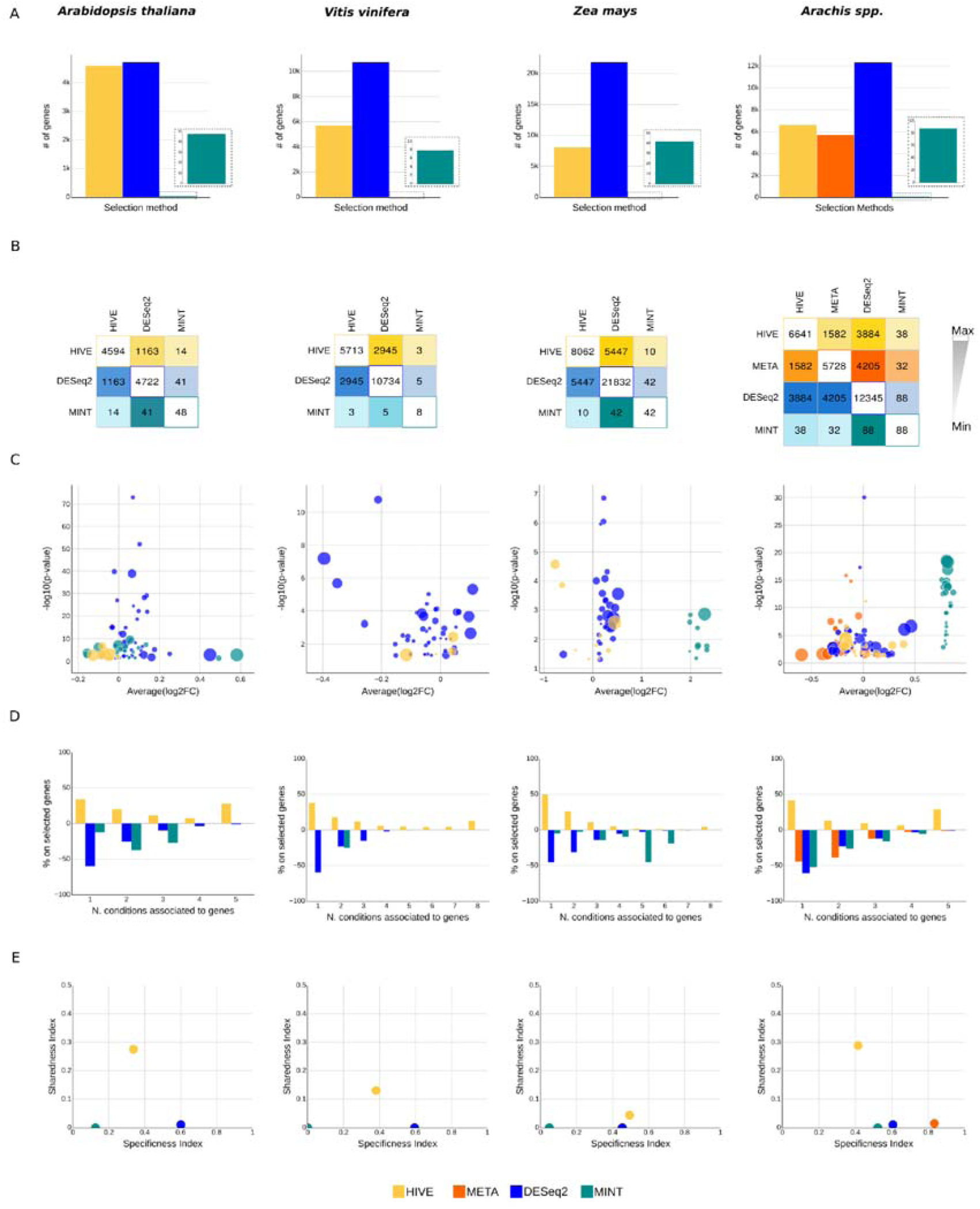
Benchmark of HIVE, DESeq2, MINT and META analysis regarding gene selection, GO term enrichment, and gene association to condition for the four RNA-seq datasets. For each biological system: A) Number of selected genes per benchmarked method. B) Pair-wise agreement in gene selection of the benchmarked methods. C) Each bubble represents an enriched GO term in the referring list (p-value <= 0.05), the size of the bubbles corresponds to the ratio of genes to which the GO term is associated and the number of coding genes for the plant of interest. On the x axis is reported the average log2FC of genes in each enriched GO term list. D) The % of selected genes associated to them by each method for each number of associated conditions. E) Relationship between the Sharedness Index and the Specificness Index of each method. (See Materials and Methods for more details on the indices calculation).

With a similar aim, we performed GO term enrichment analysis to study the biological processes in which selected genes by each method are involved (Figure 4C). To compare those results for each biological model, after selecting only significant terms with adjusted p-value lower than 0.05, we plotted the -log10(p-value) versus the average of relative expression, and the size of each point correspond to the normalized ratio between genes from the selection in each term with respect to genes in the selection compared to genes in terms compared with the background (see methods). We observe that DEseq2 selection yields the highest number of significant enriched terms as expected since it usually finds a huge number of genes when compared to the other tools. Despite, the size of the bubbles is overall smaller when compared to the other methods, thus suggesting that a lower proportion of genes in the enriched pathways was found. Surprisingly, despite the very few genes selected by MINT, the number of associated GO terms is comparable to the other methods except for grapevine model for which no terms were found enriched. However, especially for maize and peanut, we can see that the enriched terms have all a very similar average of gene expression. After manual inspection of genes belonging to the enriched terms, we found that those terms are very redundant. Overall, we observe that genes found by HIVE lead to enriched terms with a higher p-value especially when comparing with DESeq2. However, the proportion size of terms enriched in HIVE selection is higher than the other methods, thus suggesting that a high number of genes is involved in the enriched term.

An important aspect in integrative analysis is the number of signatures common to all conditions and specific to one. While the differential expression with DESeq2, or the meta-analysis specific of the peanut dataset, are calculated for each condition independently, hence this information is intrinsic with the outcome of the analysis, the genes selected by integrative methods such as HIVE or MINT are retrieved by using all conditions together. Therefore, for HIVE selection, we developed a novel strategy to associate each gene to one or more conditions which consists in calculating the amplitude of variation of each gene by measuring the distance of the expression values in each pair of conditions with the last step of the third block of HIVE (see methods, Supplementary Data 1). For MINT, since no method is provided to do this association, we applied for each conditions the same threshold on the log2FC normally used for differential expression analysis. First, we quantified the number of genes found deregulated in one condition, two conditions or more depending on the dataset for each method (Figure 4D). We observe that for the four datasets, both HIVE and DEseq2 found that around 50% of the respective selections are associated specifically to the response to one stress. While for DESeq2 the percentage of genes associated to the response to more than one condition decrease concordantly with the increase of the number of conditions, we do not observe this anti-correlated behaviour in HIVE results. As expected for HIVE we observe a consistent percentage, between 4,41% (maize dataset) and 28.89% (peanut dataser), is associated to genes found deregulated in response to all stresses, compared to a maximum of 1.04% of genes for DESeq2 selection. Regarding MINT, the results depend on the biological model. Only for the peanut dataset, it reported a number of signature specific to one stress with a comparable number respect to the other tools. Otherwise, the highest percentage for the *A. thaliana* and grapevine datasets was found for genes responding to two conditions, and to five conditions for the maize dataset. To quantitatively measure the difference among methods, we defined two novel indexes: the sharedness and the specificness which quantify the ratio of common or specific signature compared to the total number of selected signatures, respectively (see methods). We can observe in Figure 4E that DESeq2 obtained the highest values of specificness and the lowest of sharedness for peanut, grapevine, and *A. thaliana*, while for the maize datasets the specificness of HIVE is slightly higher than DESeq2. Opposite results for MINT which always achieve the lowest values of specificness for all datasets, while for the sharedness the values vary accordingly to the dataset. The sharedness and the specificness of HIVE are very similar, because it identifies a balanced ratio of common and specific signatures, compared to the other methods. Compared to other methods HIVE obtained the highest sharedness for all datasets, therefore, HIVE is able to retrieve the highest number of genes deregulated in all conditions.

Finally, we investigated the level of regulation of genes found by each method as specifically deregulated to one condition or shared. From the cumulative distributions showed in Supplementary Figure 4, we can observe that the genes selected by HIVE as specific of each condition, show the most extreme values of modulation when compared to the background, either up or down regulation, with the most significative p-value compared to the other methods and background. This result highlight that the genes found by HIVE as responsive specifically to one condition have an overall level of regulation which is stronger than the genes selected by other methods for the same category. Concerning the shared signatures, i.e. genes modulated in all conditions, we expect a global milder level of regulations compared to specific signatures. Overall, independently by the selection method, we observe that the cumulative distributions of those shared signatures are closely distributed to the background. Since in HIVE model there are no constrain on expression values, the model is able to also capture shared lowly expressed signatures missed by other methods. Those results corroborate the suitability of HIVE to analyse unpaired multi-conditions transcriptomic data, outperforming the other methods available which are not tailored for this kind of analysis.

The very similar results achieved in all datasets reinforce the robustness of HIVE findings and suggests its potential application to different biological systems.

Altogether, these results showed the advantages of performing an integrated analysis with HIVE by capturing the highest number of genes deregulated in all conditions for each dataset and the specific responding genes to one condition with the most extreme regulation levels and the achievement of better performances compared to the state-of-the-art tools.

### HIVE extracts multi-stress responding genes from the integrated analysis of multiple single-stress experiments

Among all the conditions available for each dataset, we dispose of the configuration in which the plant is subjected to two stresses contemporarily (either biotic and abiotic or two biotic agents) and to only one of the two stressors at the time (Table 2). To test the performances regarding the identification of genes responding to multiple stresses, from each dataset we extracted the lists of genes found by either HIVE or DESeq2 associated either to the combined stress or one of the two single stresses, and we compared the overlap of those lists per method. We excluded MINT from this analysis because too few genes were identified in those conditions to perform a comparative analysis. The total number of genes found in at least one of the three conditions for each model ranges from 3020 to 4011 for HIVE and from 1834 to 8856 for DESeq2 (Figure 5A, Table 2). The two extreme values for DESeq2 are due to the absence of genes associated to the *A. thaliana - Laccaria bicolor* interaction experiments and the 4303 (49% of the total) genes associated to the abiotic condition for the maize dataset. The percentages of genes found associate to both single stresses and to the combined varies from 14% to 51% (median of 34%) for HIVE versus the range of 0 to 7% (median of 3%) for DESeq2. In general, the percentage of genes associated to both single stress but not found in the combined is very low and comparable between the two analysis methods: from 1% to 6% for HIVE and from 0 to 6% for DESeq2. Regarding the percentage of genes associated specifically to the combined stress and not found either in common with one or two single stresses is lower for HIVE selection than the DESeq2 for each model.

**Figure 5.**
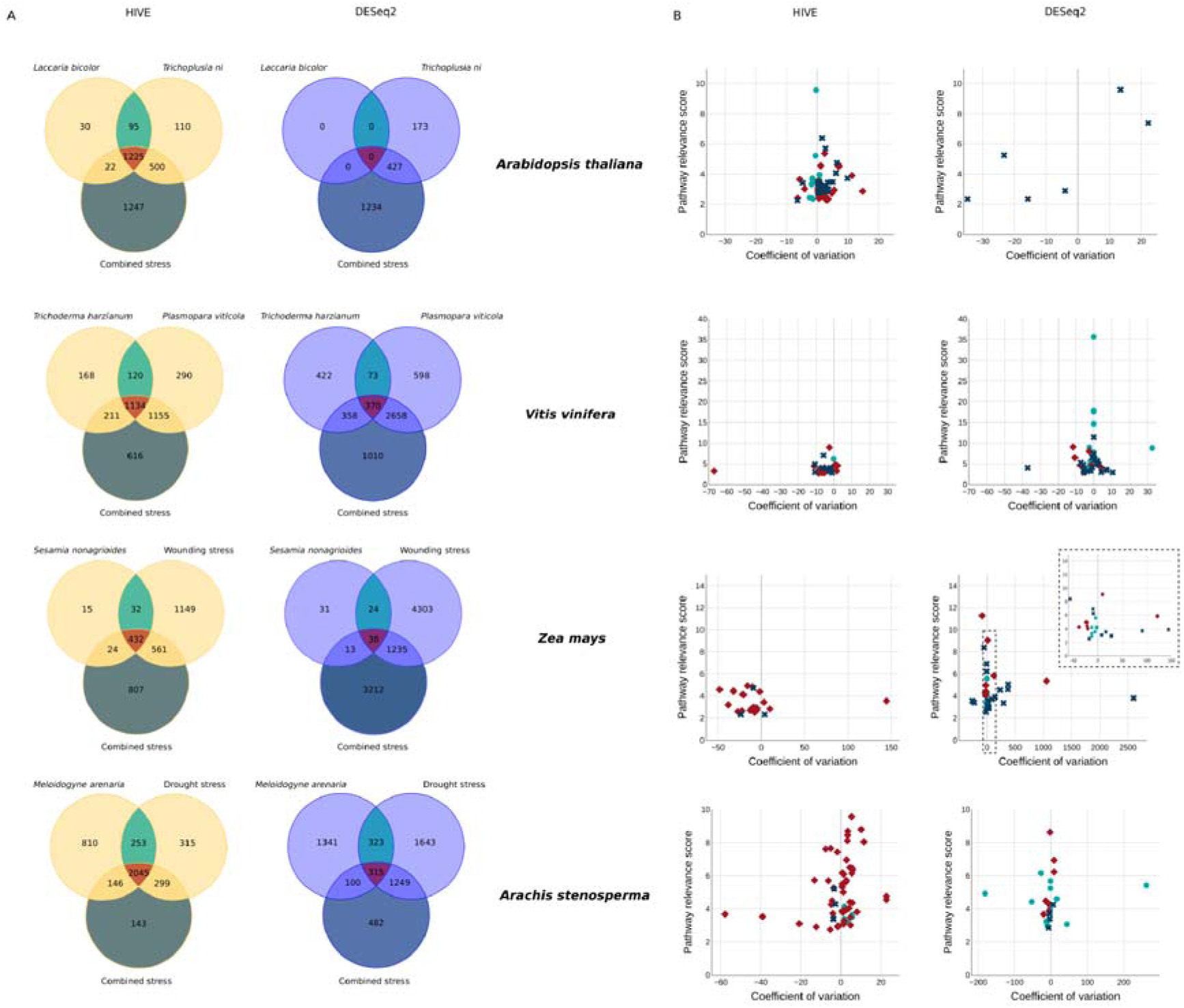
Comparison of multi-stress associated genes’ selection from HIVE and DESeq2. A) Venn diagrams representing overlaps between single stress genes and the respective combination of stresses for each plant. B) The pathway relevance for each enriched GO term in each set of genes, versus the coefficient of variation of the corresponding log2FC of the associated genes, calculated from genes in the green, red or blue portions of the Venn.

**Table 2.** Comparison of selected genes involved in two single stresses, in combined stress or an overlap of the three by both HIVE and DESeq2. The table shows the combination of stress considered for each RNA-seq dataset and elucidates the differences in multi-stress gene association in the gene lists of HIVE and DESeq2. The first column reports the considered plant dataset, the second one describe the type of stresses and the third one the details of the stresses. The forth column gives the number of selected genes from HIVE associated to at least one of the stress of interest in this analysis, the fifth column gives the % of genes on the total that are common to all stresses (two single stresses + the combined stress), the sixth gives the % of genes belonging only to the combined stress (thus not overlapping with both the single stresses) and the seventh column shows the % of the selected genes in common just between the two single stresses. Columns from eight to eleven shows the same concepts but applied to DESeq2 selection.

| Plant | Type of combined stresses | Description of combined stress | Total genes | % Common to all stresses | % Only combined stress | % Common to single stresses | Total genes | % Common to all stresses | % Only combined stress | % Common to single stresses |
| --- | --- | --- | --- | --- | --- | --- | --- | --- | --- | --- |
| *Arabidopsis thaliana* | Biotic+ Biotic | *Trichoplusia ni*+<br>*Laccaria bicolor* | 3229 | 37.94 | 38.62 | 2.94 | 1834 | - | 67.28 | - |
| *Vitis vinifera* | Biotic+ Biotic | *Trichoderma harzianum*+<br>*Plasmopara viticola* | 3694 | 30.70 | 16.68 | 3.25 | 5489 | 6.74 | 18.40 | 1.33 |
| *Zea mays* | Biotic+ Abiotic | *Sesamia nonagrioides*+<br>wounding<br>caused stress | 3020 | 14.30 | 26.72 | 1.06 | 8856 | 0.43 | 36.27 | 0.27 |
| *Arachis stenoperma* | Biotic+ Abiotic | *Meloidogyne arenaria*+<br>drought stress | 4011 | 50.98 | 3.57 | 6.31 | 5453 | 5.78 | 8.84 | 5.92 |

Overall, HIVE identified a lower number of genes which is specific to the combined stress compared to DESeq2 and most of the genes associated to the combined stress response reported by HIVE, were found also associated to the response to the two single stresses, contrarily to the DESeq2 findings.

To better understand the biological meaning of the different gene lists found by the two tools, we performed a GO term enrichment analysis on: genes found deregulated only in the combined stress, genes deregulated in response to both the two single stresses and the combined and genes found deregulated to respond to both single stress separately but not to the combined. To compare the results from the two tools, we define the “pathway relevance score” which considers the contribution of the p-values and the proportion of the number of genes found deregulated compared to the number of genes in the pathway and the number of genes in the lists (see methods for more details). Then we plotted the pathway relevance score versus the coefficient of variation, which consider the variance of gene expression levels among conditions in each GO term with respect to the average. For each plant, we can observe that the ranges of pathway relevance score are comparable between the GO term analysis results performed on each tool’s list independently (Figure 5B). Importantly, we can observe that while most of the enriched GO terms with the highest pathway relevance scores were found for the genes deregulated in the three conditions concerning HIVE, regarding DESeq2 the correspondent high values are associated to genes associated to either the combined stress or the two single stress only. Interestingly, when calculating the coefficient of variation from gene in the enriched GO terms, we found higher values from HIVE selection, compared to DESeq2. On the top of that, for each biological model, the highest number of enriched GO terms was found for genes deregulated in the three conditions for HIVE. Those results show that HIVE found a higher number of GO terms associated to genes in common between the two single stresses and the combined stress also with higher coefficient of variation compared to DESeq2 findings highlighting the efficiency of HIVE to retrieve genes with high variability across conditions compared to DESeq2 selection.

The few genes associated specifically to the combined stress and the high number of genes in common between the combined stress and both the single stresses from HIVE and their higher coefficient of variation, highlight the ability of HIVE to perform *in silico* multi-stress integration of experiments conducted on single stresses. This represent a proof-of-concept that HIVE can be used to reliably extract multi-stress signatures from experiments performed on single stresses.

### The Voronoi maps allow a global comparison of the level of sharedness – level of expression relationship across conditions

To further inspect the characteristics of the genes found by HIVE deregulated in each single stress and/or in the combined, we used the Voronoi representation. First, the genes found deregulated in either one of the two singles stress and the combined, were stratified in three levels: 1, genes exclusively deregulated in the specific condition, 2, genes deregulated in two conditions (one of the two must be the condition to which the map refers to), 3, genes deregulated in the three conditions. We refer to those as levels of sharedness. Then, for each gene in each level we extracted the level of expression in the correspondent condition. Finally, for each condition, genes were mapped accordingly to their level of sharedness-gene expression value on a x-y plot to build a Voronoi diagram. A Voronoi cell represents a volume of influence of the data point it contains, providing a direct precise measurement of the local density. The smaller the volume, the closer the genes on the Voronoi diagram, implying a high number of genes sharing properties along the 2D representation. Using this approach, we obtained three Voronoi maps, one for each condition, for each of the four plants (Figure 6).

**Figure 6.**
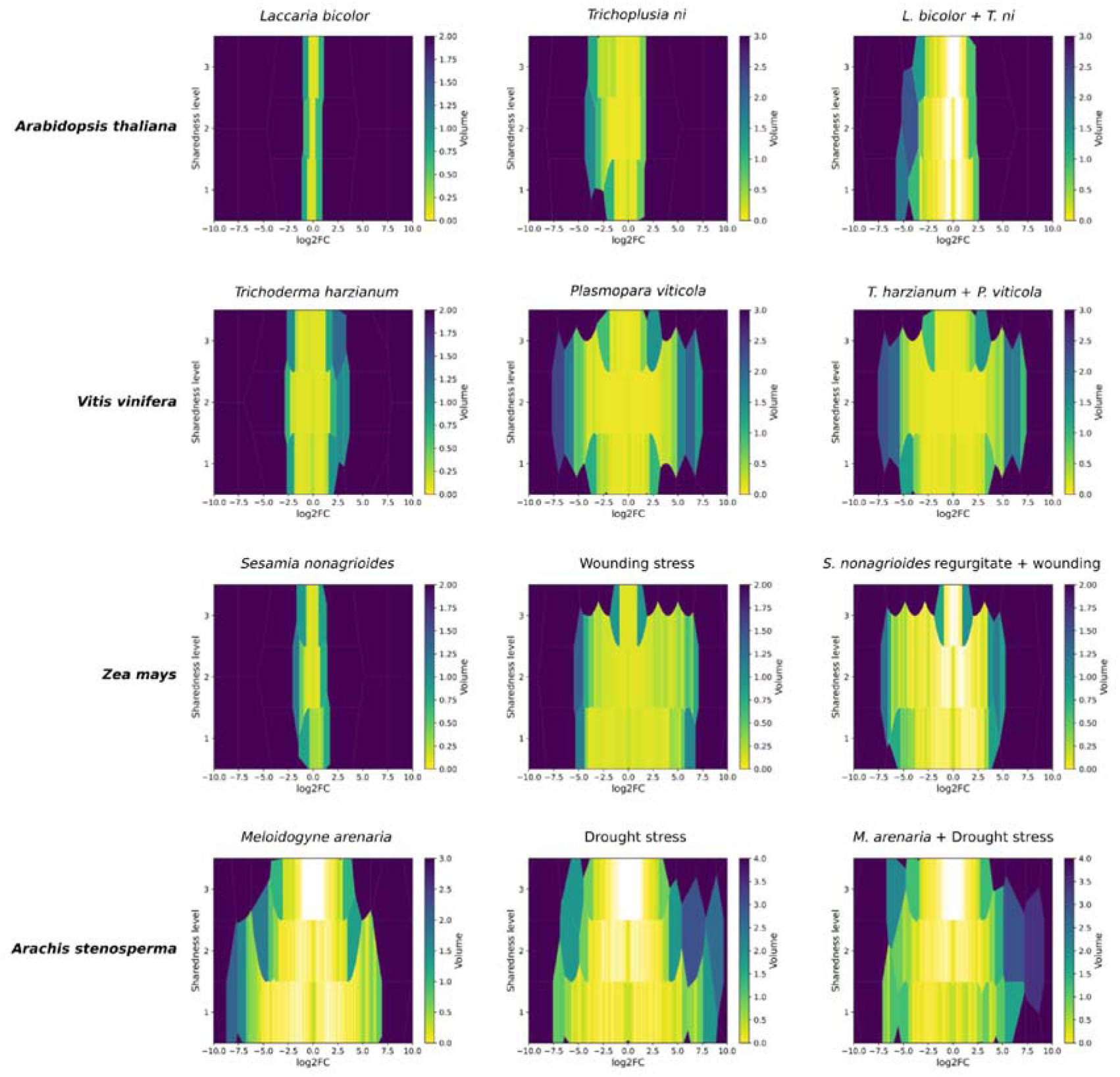
Voronoi maps representing gene expression in different sharedness levels across conditions. The first two Voronoi maps of each raw, show the expression of genes associated to the single stresses, while the last one represents the expression level of genes associated to the combined stress, for each biological system. Genes were mapped accordingly to their level of sharedness-gene expression value on a x-y plot, each gene were then enclosed in a Voronoi cell, the volume of each cell is represented by a colour scale, accordingly to each plot.

In level 1 only genes associated to each specific condition are reported, thus genes in this level are only present in the specific map representing the stress in which the gene was found exclusively deregulated. On the other hand, genes in level 2 are present in two maps since those genes were found to be responding to two conditions. By definition, genes associated to the response to the three conditions are in level 3 and therefore are represented in the three maps. The color scale depends on the volume of each Voronoi cell, each containing only one gene, yielding an index of the density of the different regions in the Voronoi diagram. The more similar is the density of volumes for each level in the maps, the more similar is the overall response of the compared stresses. The advantage of this representation is that it allows to easily compare the overall response to different stresses affecting the same plants but also among different plants. We can observe that for maize, grapevine and *A. thaliana*, one stress yields a lower response that the other, while for peanut both stresses trigger similar amplitude of response. Accordingly, we observe that the density of Voronoi cells in level 3 is lower for those three plants compared to peanut. Interestingly, for maize and grapevine the Voronoi diagram of the combined stress in very similar to the map obtained for the stress condition with the highest effect, suggesting that the stress with the milder effect do not trigger additional responses when also the second stress, with a stronger effect also alone, is present. This is consistent with the conditions studied in those models. The milder stress in grapevine is due to the fungi *Trichoderma harzianum* while a stronger effect is observed in presence of *Plasmopara viticola*. In the original publication they showed similar results in which *T. harzianum* enhance grapevine response to *P. viticola* but do not induce a strong response when present as only stressor (79). For maize we observe, consistently with previous findings (80), that the mechanical stress due to wounding induce a long-term strong response opposite to the presence of only or combined of *Sesamia nonagrioides*. Although for the *A. thaliana* experiments there is a condition triggering only a very mild response as for maize and grapevine, the Voronoi diagram for the combined stress is not similar to the strong stress associated map. This might suggest that even if the presence of *Laccaria bicolor* alone seems not to affect the gene expression program dramatically, when *Trichoplusia ni* is infecting the plant, the global effect of the multifactorial stress is different compared to each stress separately. Also for this model, this result was expected since the presence of *Laccaria bicolor* enhance the plant response to *Trichoplusia ni* (81). For the peanut conditions, we observe that the Voronoi diagram for each stress separately have a higher density of cells especially in level 1 and 2 and higher log2FC range than the map for the combined stress. In level 3, we can see similar distribution of volumes denser in the center of the plots, implying not only similar level of regulation in the three conditions, but overall with milder log2FC than specific responding genes (level 2 and 1). This suggests that the peanut plant subjected to both stresses have similar response to when under only one of the two at the time.

To quantify the differences in the cells’ density, we used their volumes and calculated the volumes distributions of cells by ranges of log2FC for each level and compared those distribution for each stress (Supplementary Figure 5). The volumes distributions recapitulate the findings qualitatively observed from the Voronoi diagrams. Strikingly, while the volumes distributions of cells from wounding stress and P. *viticola* for maize and grapevine respectively and each respective combined show no significant difference in level 2, they are different from *S. nonagrioides* infection and *T. harzianum* infection distributions. For *A. thaliana*, the volumes distributions are significantly different from the other two stresses in all levels. Finally, for peanut the difference is mainly observed in level 1, with distributions of volumes from the combined stress map higher than the other two stresses singularly, while they are more similar in the other two levels.

In conclusion, the Voronoi diagrams yield an easy visualization of the relationship between the levels of sharedness and the level of expressions. The inspection of those maps allowed to identify which combination of sharedness-expression are more (or less) frequent in each condition. Therefore, the Voronoi representation allows a global comparison of the sharedness-expression relationship across conditions and therefore to identify conditions with similar (or different) patterns. This analysis highlights that the patterns of sharedness-expression found by HIVE on the single stresses are comparable to the combined, therefore strengthening the ability of HIVE to perform in silico multi-factorial stress integration from single stress data.

### Latent features extracted from the peanut dataset are associated with phenotypic characteristics

In the previous analyses we showed that the HIVE achieved the best performances in reducing the batch effects, selected both common and specific signatures in response to multiple and single stresses respectively and identified the highest number of genes responding to the combined stress and found also responding to both single stresses, on the four datasets. To deeply investigate the biological insight found by performing *in silico* multifactorial stress with HIVE, we decided to focus on one dataset. We chose the peanut dataset to inspect whether the integrated analysis using HIVE could retrieve novel signatures beyond the original findings in Mota et. al., 2021 (37), important to decipher the arachis response to biotic and/or abiotic stress.

First, we explored whether the latent features encoded in the latent space can capture patterns associated with the different phenotypic characteristics in the dataset, by using the point-biserial correlation and then Kruskal-Wallis followed by the Dunn’s test. Regarding the peanut dataset, first we studied the two main characteristics, namely the type of stress (biotic or abiotic) and the type of plant species (*A. duranensis* or *A. stenosperma*). By using the point-biserial correlation, we calculated the most significant latent features correlating either with the type of stress or the species (see methods). Among the 80 latent features, we found 19 features correlated with plant species and 16 with biotic/abiotic stress (Figure 7A). Interestingly, three of them were found significantly correlated to both characteristics. Then we plotted the two most significative latent features associated with each phenotypic characteristic and we observed samples grouping accordingly (Figure 7B and 7C). The Kruskal-Wallis followed by the post-hoc Dunn test, was then used to compare pair-wisely the conditions and counting how many latent features are associated to the discrimination of each pair of experimental condition (Figure 7D). *A. duranensis* biotic and *A. stenosperma* drought showed the highest number of latent features (30 latent features) associated to their difference followed by *A. duranensis* biotic and *A. stenosperma* cross and *A. duranensis* biotic and *A. stenosperma* biotic (29 latent features and 28 latent features respectively) as expected therefore confirming the validity of the latent features to resume the multiple phenotypes of the datasets. Interestingly, only one latent component was found associated with differentiating *A. stenosperma* drought and *A. stenosperma* cross, which was not expected by the results of the meta-analysis performed in the original publication.

**Figure 7.**
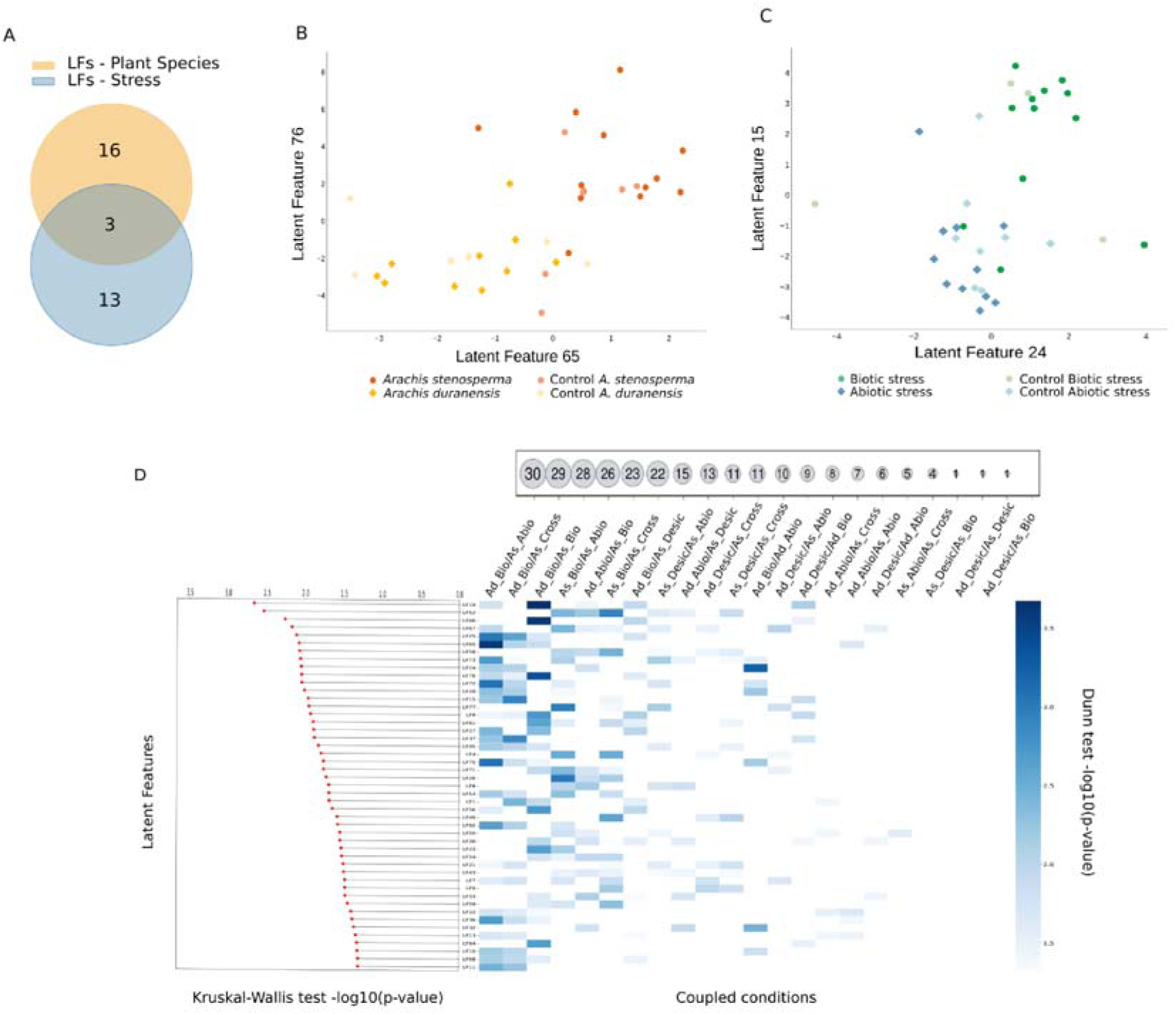
Latent features are associated with phenotypic characteristics. A) Number of latent features selected by the point biserial correlation (p-value < 0.01) as the most correlated either with the type of stress (yellow) or plant species (light blue). Distribution of samples in the space of the top two correlated latent features with plant species (B) or with type of stress (C). The condition is indicated in the correspondent legend. D) Kruskal-Wallis test (on the left) and post-hoc Dunn test (on the right), application to pairs of conditions for each selected latent feature (“Desic” stands for desiccation stress). On top-right is represented the number of differing latent features accordingly to the Dunn test results.

Overall, these results show not only that the latent features reduce the batch effect but also that they capture the phenotypic characteristics of the dataset.

### The integrative analysis with HIVE allowed to identify the genes involved in the specific or common response to biotic and abiotic stresses in arachis spp

We inspected the functional pathways associated to common and specific signatures to understand which biological processes are commonly modulated in response to both stresses and in both species and which are very specific of only one condition.

To analyse the common signatures, we selected the 1919 genes modulated in all conditions and performed GO term enrichment analysis and reported for each condition the average expression values of genes belonging in significantly enriched terms (Figure 8A and Supplementary Figure 6). Interestingly by clustering the conditions based on those average expression profiles, we observe that the clusters correspond to the similarity of the phenotypes. Strikingly, very few enriched terms associated to the selected common genes are modulated similarly in all condition, but most are modulated depending on the stress and/or genotype. This result suggests that despite the same genes and terms are involved in response to both biotic and abiotic stress in both species, their activation or repression can be specific of each condition. For instance, while genes involved in translation and macromolecule biosynthetic process show the tendency to be upregulated in all conditions, regulation of gene expression and nucleobase-containing compound metabolic process show overall down-regulation in almost all condition while regulation of metabolic process is down-regulated only in one time point of biotic stress for each plant genotype. Importantly, despite those genes were found to be deregulated in all conditions, we observe a very similar pattern of expression between abiotic and cross-stress and biotic (especially 6 days) and cross-stress, reinforcing the ability of HIVE to capture shared signatures from single stress conditions.

**Figure 8.**
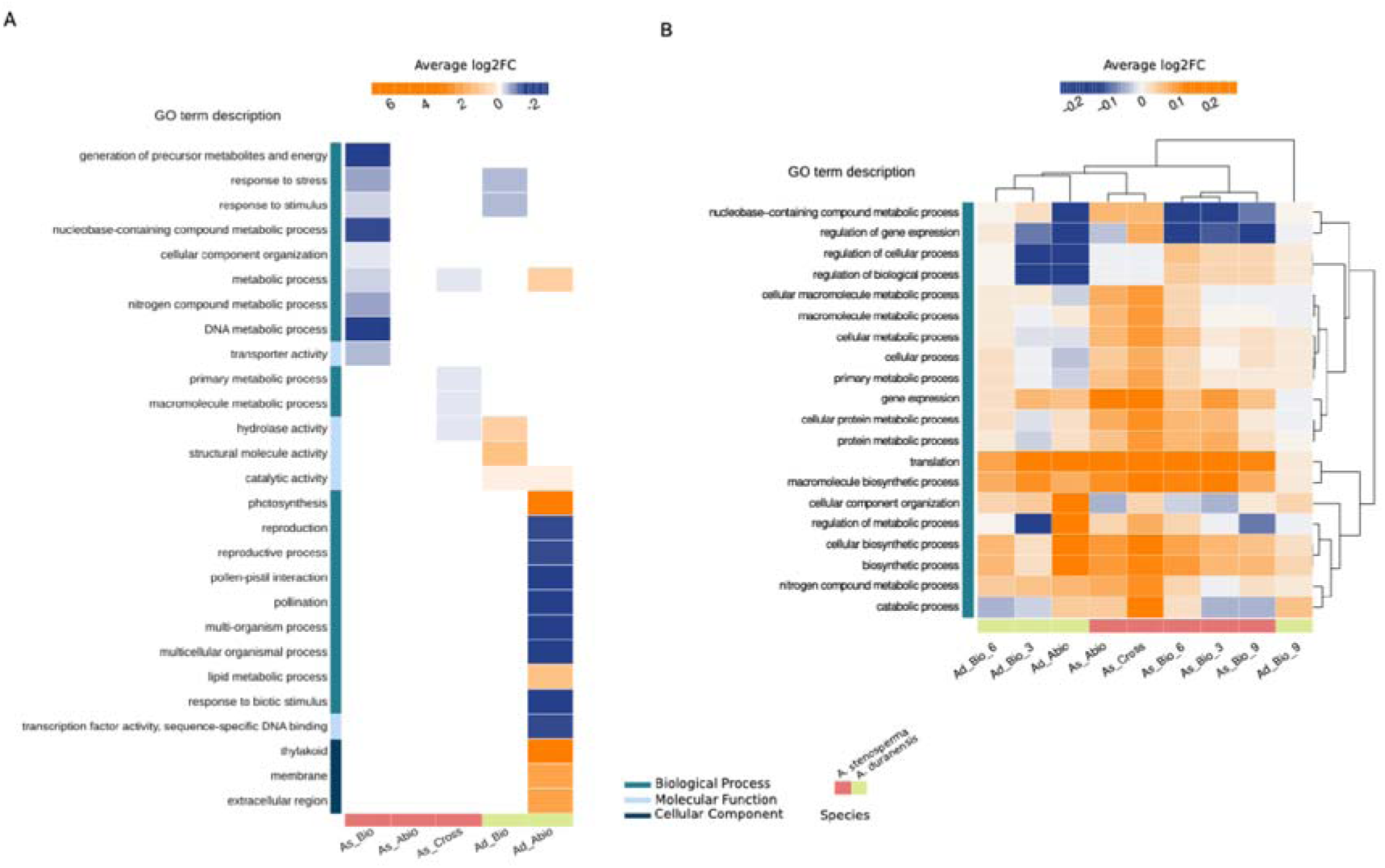
Functional analysis of genes associated to common or specific conditions. Average log2FC of genes in enriched GO terms (p-value <= 0.05) from HIVE selection A) for genes associated to specific conditions or B) associated to all conditions.

Then, we inspected the specific signatures by performing the same analysis on the genes that were found to be modulated specifically in only one condition (Figure 8B). Strikingly, although the genes selected for each condition are different, we can observe that the same terms are found enriched in multiple conditions, suggesting that the same pathway is modulated in response to both stresses and species but different genes are involved. For example, the term “response to stress” is enriched in response to biotic stress in both genotypes, despite 23 genes are modulated in *A. stenosperma* and 9 in *A. duranensis*. Interestingly, although the selected genes are different, the global regulation of the pathway is similar. Other terms are specific of one unique condition, such as DNA metabolic processes, whose correspondent 8 genes are downregulated in *A. stenosperma* during biotic stress showing a down-regulated average; or thylakoid cell compartment which shows an average up-regulation of 36 genes only in *A. duranensis* abiotic. Of note, we observe very few terms associated to specific genes responding only to one stress in *A. stenosperma*, because most of the signatures associated to one stress are also associated to the cross-stress as already showed by previous analysis (37).

Altogether these results show the importance of performing the integrated analysis to retrieve both common and specific signatures in response to the multiple stresses studied and the validity of HIVE findings.

### HIVE identifies genes involved in the main hormone signalling pathways, kinases, and resistance in response to stress

To further characterize whether HIVE can capture genes involved in plant responses to stresses, we inspected the 6641 genes found by our integromics approach searching for genes involved in the main phytohormones signalling pathways, kinases and resistance genes.

93 genes were identified belonging to seven hormone signalling pathways, three directly connected with abiotic/biotic stress response (abscisic acid, jasmonate and ethylene) and four growth-promoting hormones (gibberellins, auxins, cytokinins and brassinosteroid). Considering the pivotal importance of plant hormones in mediating the regulation of stress responses, we focused on studying the expression profiles of these genes using a hierarchical clustering approach. Overall, we observe that these genes are activated or inhibited with very different patters depending on the stress and the plant species (Supplementary Figure 7A). Nevertheless, genes involved in Jasmonate are mainly activated in *A. stenosperma* in response to nematode infection probably because of the known role of this signalling pathway in biotic stress response for plant protection (82–84). Interestingly, the expression values for those genes upon cross-stress are slightly lower than during biotic stress only, while mild downregulation is observed during abiotic stress only for *A. stenosperma*. Brassinosteroids are activated/inhibited in a stress and plant species specific manner. The two growth hormones Gibberellins and Cytokines related genes show similar expression profiles including while some genes are activated in both stresses and plant species, others show a more specific pattern.

Interestingly, genes in the Ethylene pathway are strongly activated in *A. duranensis* and mildly in *A. stenosperma* during RKN infection. Finally, genes in the Abscisic and Auxin pathways have a very heterogeneous patter of expression with an overall activation in response to drought stress mainly for Abscisic related genes.

Previous analysis of the separated conditions showed that the jasmonic acid pathway was prevalently induced during biotic interaction, while Abscisic acid upon abiotic stress (37). The integrated analysis allowed to identify other important hormone pathways involved in defence response and their different modulation depending on the plant species.

Plants have evolved a variety of mechanisms to cope with environmental changes, among which kinases play an important role in regulating both plant development and stress responses (85–87). Plant kinases are originally sensors for a variety of signals like phytohormones and small peptides, facilitating intercellular communication. Among HIVE-selected genes, we have found 264 genes coding for kinases. By inspecting their transcriptional profiles, we identified 8 clusters (Supplementary Figure 7B). Three clusters (A, E, G) group kinases induced in a stress and plant species-specific fashion (abiotic - *A. duranensis*, biotic and cross stress - *A. stenosperma*, abiotic and cross-stress - *A. stenosperma*, respectively). Kinases in cluster D are activated in response to biotic stress in both plant species and cross-stress for *A. stenosperma*. The remaining four clusters bear genes showing modulated activation in multiple stresses and plant species at the same time.

Other key players of plant immunity are resistance genes (R-genes) (85). They code for the R-proteins, which have a main role in recognizing secreted pathogen-specific proteins (effectors) and activate a powerful, pathogen-specific immune response, which often includes transcriptional re-programming (88). Most of the R genes in plants encode nucleotide-binding site leucine-rich repeat (NBS-LRR) proteins (89). This large family is encoded by hundreds of diverse genes per genome and can be subdivided into the functionally distinct TIR-domain-containing (TNL) and CC-domain-containing (CNL) subfamilies (90). Usually, those genes have very low expression profiles making them hard to find with classical analysis methods (91). HIVE identified 79 NBS-LRR genes whose expression profiles are reported in Figure 6C. Remarkably, the meta-analysis found 55 NBS-LRR, reinforcing the utility to perform integrated analysis using HIVE. We mainly observe similar patterns as for the kinases: either the genes induced upon specific stress-plant species or heterogeneous patterns showing modulations in multiple conditions, the latter were not previously found by non-integrated analysis methods (35). Of note, seven of them were previously validated by qRT-PCR experiments (35), reinforcing the reliability of the results found by HIVE.

In conclusion, by integrating experiments on different plant species each affected by one stress, HIVE can capture genes triggered in the different conditions linked to main hormone signalling pathways, kinases and R-genes in response to stress.

### The integrated analysis allowed to infer the complex gene regulatory network induced by biotic and/or abiotic stress

Plant responses to stress are controlled by a complex gene regulatory network (GNR). Therefore, we focused to study the transcription factors (TF) – targets relationship. Then we collected TF information from PlantRegMap and we found 69 TF among the HIVE selected genes distributed in 21 families (Figure 9A). We inferred the GNRs by considering those 69 TF as the “regulators” and the rest of the 6287 genes that are not TFs as targets. By using the GENIE3 model (68) (see methods), we inferred 2214 connections. Then we filtered out the pairs not in agreement with the prediction from PlantRegMap, obtaining 22 TFs and 127 interactions (Figure 9B). Interestingly, by inspected those pairs we have found two TFs regulating genes involved in phytohormone pathways. The bZIP TF Aradu.C0RFP and the Ethylene-related gene Aradu.MA5U7, with anticorrelated expression profiles (Supplementary Figure 8A) and the ERF TF Aradu.K41I0 with the gene Aradu.1C9UI involved in the Jasmonic acid pathway, showing correlated expression profiles (Supplementary Figure 8B). We performed the same analysis on the genes selected by the meta-analysis. Among the 379 TFs in the meta-analysis, 76 are also present in PlantRegMap, yielding 5349 genes as putative targets. Applying the same method, we selected 2919 pairs. By filtering out the pairs not in agreement with PlantRegMap we obtained 23 TFs and 86 interactions (Figure 9C), 32% less than the pairs found with HIVE genes selection. These results suggest that HIVE captures more reliable signatures to infer GRN.

**Figure 9.**
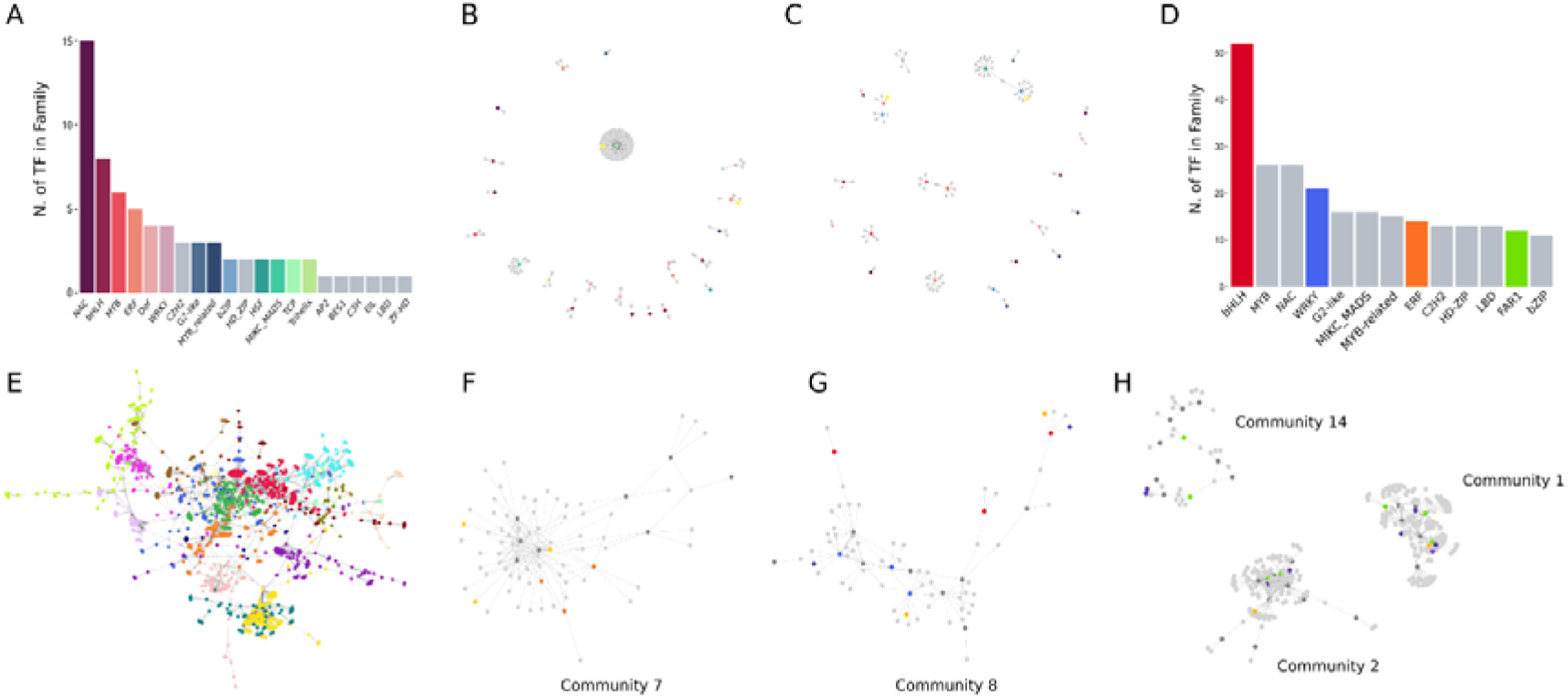
The integrated analysis allowed to infer the complex gene regulatory network induced upon biotic and/or abiotic stress. A) Number of transcription factors from HIVE selection with predicted binding motif in PlantRegMap. B) GRN inferred by GENIE3 imposing as “regulators”, the TF found by HIVE and PlantRegMap, and as “targets”, genes that are not TFs (after having removed all pairs not in PlantRegMap). Nodes are coloured accordingly to A. C) same as B but with gene selection from the meta-analysis. D) Number of transcription factors in families with at least 10 representatives in HIVE selection. E) The largest connected network from the GRN inferred by GENIE3 imposing as “regulators”, the TF found by HIVE and not in PlantRegMap, and as “targets”, genes that are not TFs. Colours correspond to the 19 communities found by Leiden algorithm. F) Zoom-in into community n. 7, orange nodes correspond to transcription factors of family ERF. G) Zoom-in into community n. 8, bright blue nodes correspond to transcription factors of family WRKY, and red nodes to transcription factors of family bHLH. H) Zoom-in into communities n. 14, 1, and 2 bright green nodes correspond to transcription factors of family FAR1. In all network figures dark grey nodes represent all the other transcription factors in the network not belonging to the family/families of interest, dark yellow nodes correspond to genes associated with phytohormones, violet nodes indicate NBS-LRR genes and all the light grey nodes are other categories of regulated genes.

To reconstruct a more general regulome, we extracted all the TFs among the genes selected by HIVE analysis independently by their presence in PlantRegMap (Supplementary Table 4). Among the families with more than 10 TFs found by our analysis, we observe families with known pivotal roles in development, plant growth and stress responses (Figure 9D) (68, 92). Then we inferred another network using the same strategy as before, from the 354 TFs in HIVE selection, we removed the 69 used in the previous network for a total of 285 TFs as “regulators” and the 6287 genes as “targets”. By selecting the top 2500 pairs, we obtained one large connected community and several isolated interactions including 224 TFs and 1410 targets (Supplementary Figure 9). We focused on the main network for further investigations, and we applied the Leiden algorithm to find communities of genes densely connected. Leiden community detection is a fast and accurate method that allows us to find well-connected and refined partitions of the network. We obtained 19 communities as shown in Figure 9E with variable number of TFs and families distribution (Supplementary Table 5). We first inspected community 7 which shows the highest percentage of phytohormones related genes with respect to the total number of genes in the community (Supplementary Table 6), involved prevalently in the Jasmonic and Abscisic acid pathways (Figure 9F). Interestingly this community also presents a high percentage of TFs belonging to the Ethylene Responsive Factors (ERF) family. The expression pattern of these genes and TFs are very similar among each other, all showing high activation in *A. stenosperma* under RKN infection but different patterns in the other conditions. Two TFs (Aradu.LD7BF and Aradu.3P75R) highly repressed in *A. duranensis* in the earliest time point of *M. Arenaria* infection, two hormone gene repressed similar to the previous but one (Aradu.EQ5J6) also in drought stress Aradu.WB9GS, while Aradu.E2TII (TF) and Aradu.XW8AZ (hormone gene) strongly repressed in *A. duranenesis* in drought condition (Supplementary Figure 10A). The crosstalk among these three hormone pathways was already suggested in the previous publication related to those data. Despite their enrichment we do not observe a direct interaction between the ERF TFs and the hormones genes involved in jasmonic and abscisic acid pathways suggesting that other intermediary TFs and genes intervene to regulate this complex crosstalk therefore providing novel potential key interactions to further inspect. The second most enriched communities in genes related to phytohormones pathways are community 8 and 2. In community 8 there are also TFs of WRKY and bHLH families and stress-response related genes. Therefore, we screened the community for NBS-LRR genes (Supplementary Table 7) and we found two as direct targets of TFs belonging to the two enriched families: Aradu.180Q6 (WRKY TF) – Aradu.NP598 (NBS-LRR) and Aradu.X5F2F (bHLH TF) – Aradu.HLR71 (NBS-LRR) (Figure 9G). By inspecting the expression profiles Aradu.NP598 (NBS-LRR) shows a different patter compared to the other selected genes in the community and strong repression in *A. stenosperma* after 6 and 9 hours of *M. arenaria* infection, while Aradu.HLR71 (NBS-LRR) show mild activation in *A. stenosperma* independently by the stress with similar expression profile to the putative TF regulator (Aradu.X5F2F) (Supplementary Figure 10B). In community 2, the most represented family of TF is the far-red-impaired response 1 (FAR1), this family is present with the highest percentage also in community 1 and 14. Intriguingly, community 1 bears the highest percentage of NBS-LRR genes. Therefore, we decided to analyse those three communities altogether (Figure 9H). This family of TFs was shown to be implicated into pod development in *A. hypogaea* (93). Furthermore, we have observed high percentage of gibberellin related genes, whose modulation was shown to be important for controlling pod size (94). In the three communities we observe NBS-LRR genes either as direct targets or as secondary targets of FAR1 TFs, probably suggesting a possible regulatory role for this family in R-gene mediated Arachis response to stress. Among this last kind of interactions, in community 14 we remarked that two NBS-LRR genes (Aradu.RT9LH and Aradu.G29LA) are targeted by the same TF (Aradu.VL3IZ) of the WRKY family that is found to be a target of a FAR1 TF (Aradu.1I5WA). The expression profiles of the FAR1 TF and the two NBS-LRR is opposite in all condition, with Aradu.G29LA showing higher differences compared with Aradu.RT9LH, suggesting an opposite activity mediated by the intermediary WRKY TF (Supplementary Figure 10C). Overall, the integrated analysis with HIVE allowed the selection of genes with pivotal roles in stress responses and joined to GENIE3 model, to infer a comprehensive GNR to better understand the crosstalk among the different strategies used by plants to respond to diverse stresses.

### Plant functional validation of NBS-LRR genes found by HIVE

Since the main scope of the development of HIVE is to improve the identification of signatures responding to multi-stress conditions, we decided to choose candidates genes for experimental validation which were found to be modulated in response to multiple conditions. Those candidates might confer a broader resistance either in multiple plant genotype or multiple stresses. Therefore, among the NBS-LRR genes found in the inferred regulome, we selected two genes showing modulation in multiple conditions to be validated *in planta*. The first NBS-LRR gene, *AsTIR19* (Aradu.G29LA), was identified in both the highly resistant species *A. stenosperma* and in the moderate resistant *A. duranensis* in response to the infection of the RKN *M. arenaria*. This gene was found in community 14, among the highest activation in *A. duranensis*. The second NBS-LRR gene, *AsHLR* (Aradu.HLR71) from community 8, was mildly activated in *A. stenosperma* in the response to both biotic and abiotic stresses. Importantly, this gene was not identified by the analysis performed in the previous publication using the meta-analysis as data analysis technique. Since this gene respond to both biotic and abiotic stress separately and in the cross-stress, it was missed by the meta-analysis which mainly identify signatures responding to one single stress as showed by previous paragraphs. For *in planta* validation of these NLR genes, we produced six *A. thaliana* overexpressing lines (OE) for each gene, and challenged them with the RKN *M. incognita.* For *AsHLR*, although all six OE lines showed a reduction in nematode infection in relation to WT, only HLR-OE 4.1 line showed to be significant (p<0.05), with an average reduction of 28% in the number of galls/g of root (Supplementary Figure 11A). The overexpression of *AsTIR19* in Arabidopsis showed a significant reduction of RKN infection in all six OE lines tested (Supplementary Figure 11B), with an average reduction of 52.8% in the number of galls/g of root. These results suggest that, both genes, alone or in association with other defence genes, have the potential to increase *M. incognita* resistance in transgenic plants. Therefore, the results obtained here, which combines the identification of candidate genes by HIVE followed by *in planta* validation, have the potential to be relevant to other plants affected by this pathogen.

The *in planta* validation of those two genes and mainly of *AsHLR,* which was not identified by the previous study performed using the meta-analysis, confer more robustness and reliability to HIVE findings and confirm its ability to find multi-stress responding genes from single-stress studies.

## Conclusions

In this paper, we presented HIVE, a novel method to integrate and analyse single stress unpaired transcriptomics data to identify multi-stress responding signatures. HIVE is an unsupervised model that uses a variational autoencoder to remove the batch effects by creating a compressed representation of the original data called latent space. This is a novel representation that captures the salient patterns to describe the experimental phenotype. With the aim of finding genes with expression profile characteristics of the studied phenotype, we developed a strategy to find genes explaining the latent features by using a random forest regression coupled with the SHAP explainer. This procedure decomposes the problem to infer the relationship between a set of N latent features and G genes into a set of N regression problems. Although variational autoencoders have already been extensively applied to analyse omics and multi-omics data, HIVE is the first model adapted for unpaired multi-conditions transcriptomics data. Furthermore, autoencoders and variants have mainly been employed for sample classification in a supervised fashion (14, 17–21). Sample classification is not among the tasks of this implementation and our model is unsupervised to capture the data structure without imposing constraints. Indeed, we showed that our unsupervised model performs better than the supervised alternative MINT (31) in the case of multiple conditions each from a different experiment and with few replicates for each condition. The main advantage of the integrated analysis with HIVE is the possibility of finding genes modulated in multiple conditions without performing several comparisons that usually yield a very poor agreement, as showed here by using DESeq2 or the meta-analysis (29, 78). The possible limitations in the performances of MINT could be due to the requirement of having all conditions sampled in all experiments to integrate, which is not the case in this study, since we are integrating multiple conditions, each coming from one correspondent different experiment. Therefore, the complex experimental design mixing multiple conditions, making also difficult to choose the classes to perform the supervised classification included in MINT and the limited number of replicates of some classes, could have affected the performances of MINT and reinforcing the need to develop a new tool to analyse more complex experimental designs.

HIVE is particularly appropriate in the field of plant biology. Indeed plants, as sessile organisms, need to continuously adapt to the changing environment and various stresses occurring altogether by reprogramming gene expression profiles. Characterizing how the regulation of gene expression is modulated in response to multiple stresses is pivotal to understand how plants respond to the biological environment in a more realistic framework. We performed extensive benchmark against other state-of-the art tools to test the ability of HIVE to reduce the batch effect, on one side and to extract biologically relevant genes, on the other side, using the eight different multi-stress integrated datasets.

Through several analysis we showed that HIVE is the only tool to identify both multi-stress and single stress responding genes. Specifically, we compared HIVE findings from the multifactorial stress experiments in which the plant is subjected to two stresses contemporarily (either biotic and abiotic or two different biotic agents) and from the plant subjected to only one of the two at the time. We showed that HIVE found only few genes associated only to the combined stress while a high number of genes in common between the combined stress and both the single stresses were retrieved, compared to the other tools. This represent a proof-of-concept that HIVE can be used to reliably extract multi-stress signatures from experiments performed on single stresses and to perform *in silico* multi-stress integration from experiments conducted on single stresses. To further inspect the characteristics of the genes found by HIVE deregulated in each single stress and/or in the combined, we used the Voronoi representation. This representation allows a global comparison of the sharedness-expression level relationship across conditions and therefore to compare the patterns across conditions. We found that the patterns of sharedness-expression from HIVE Voronoi on the single stresses are comparable to the combined, therefore strengthening the ability of HIVE to perform *in silico* multi-factorial stress integration from single stress data.

The selected case study to deeply investigate HIVE results is a dataset composed of six distinct transcriptomic experiments of two wild *Arachis* plants during root-knot nematode *Meloidogyne arenaria* infection and/or drought stress. The availability of both single stress (biotic / abiotic) separately and together (biotic and abiotic at the same time) makes this dataset the ideal showcase to illustrate the functionalities of HIVE. Specifically, this dataset allowed to show that signatures found by HIVE responding to each stress separately, were also found in the cross-stress. The other analysis methods showed lower agreement than HIVE of signatures responding to single stress and found also in the cross-stress. This result is pivotal to highlight that HIVE can provide *in silico* single-stresses integration to find multi-stress signatures.

Overall HIVE selected several genes implicated in key molecular processes involved in plant response to stress. Furthermore, by studying the pattern of expression of these genes we showed how these pathways are modulated depending on the different phenotypes. Indeed, HIVE selected genes that are exclusively activated or repressed upon one stress in a particular plant species but also more complex patterns such as genes modulated differently in the studied conditions. These results can help to better understand plant response to different stresses constituting a useful resource of potential candidate genes for engineering a broader stress tolerance. We also showed that by applying the network inference model GENIE3 (68) on genes selected by HIVE we were able to reconstruct a comprehensive interactome of *Arachis* to respond to *M. arenaria* infection and drought stress. We showed that the candidates provided by HIVE, enabled to reconstruct a network with more reliable connections than by using genes selected by the meta-analysis. The study of the inferred network allowed to better understand the crosstalk in response to biotic and abiotic stress conditions. Finally, the *in planta* validation of two NBS-LRR, one of them not identified by the previous study, both involved in multi-stress responses, yield confirmation of the reliability of HIVE results and the utility to find multi-stress responding signature from single stress experiments missed by other analysis models.

Overall, HIVE is a valuable tool to study jointly single-conditions transcriptomics data for the identification of plant multi-stress response signatures. We showed that HIVE can be applied on any phytopathosystems and will provide improvements in our understanding of the mechanisms set up by the plant to respond to simultaneously multiple stresses and allow to engineer durable and broader stress resistance.

## Acknowledgments

This work was supported by the French government, through the UCA JEDI Investments in the Future project managed by the National Research Agency (ANR) under reference number ANR-15-IDEX-01. HS was funded by a joint project of the Province of Bozen-Bolzano and the Austrian Science Fund FWF.

## Authors’ contributions

GC: methodology, software, validation, investigation, writing original draft, visualization; SM: methodology, software, formal analysis; APZM: resources, data curation, writing review & editing, MV: software; HS: supervision, writing review & editing; ACMB: investigation, resources; PMG: investigation, resources, writing review & editing; SB: conceptualization, methodology, validation, writing original draft, supervision, project administration.

## References

1. Großkinsky, D.K., Syaifullah, S.J. and Roitsch, T. (2018) Integration of multi-omics techniques and physiological phenotyping within a holistic phenomics approach to study senescence in model and crop plants. J Exp Bot, 69, 825–844.

2. Bolger, M.E., Weisshaar, B., Scholz, U., Stein, N., Usadel, B. and Mayer, K.F.X. (2014) Plant genome sequencing - applications for crop improvement. Curr Opin Biotechnol, 26, 31–37.

3. Yang, Y., Saand, M.A., Huang, L., Abdelaal, W.B., Zhang, J., Wu, Y., Li, J., Sirohi, M.H. and Wang, F. (2021) Applications of Multi-Omics Technologies for Crop Improvement. Frontiers in Plant Science, 12.

4. Sugimoto, K., Xu, L., Paszkowski, U. and Hayashi, M. (2018) Multifaceted Cellular Reprogramming at the Crossroads Between Plant Development and Biotic Interactions. Plant Cell Physiol, 59, 651–655.

5. Asai, S. and Shirasu, K. (2015) Plant cells under siege: plant immune system versus pathogen effectors. Curr Opin Plant Biol, 28, 1–8.

6. Zhang, H., Zhu, J., Gong, Z. and Zhu, J.-K. (2022) Abiotic stress responses in plants. Nat Rev Genet, 23, 104–119.

7. Zhang, H., Zhao, Y. and Zhu, J.-K. (2020) Thriving under Stress: How Plants Balance Growth and the Stress Response. Dev Cell, 55, 529–543.

8. Zhu, J.-K. (2016) Abiotic Stress Signaling and Responses in Plants. Cell, 167, 313–324.

9. Mohammadi-Shemirani, P., Sood, T. and Paré, G. (2023) From ‘Omics to Multi-omics Technologies: the Discovery of Novel Causal Mediators. Curr Atheroscler Rep, 25, 55– 65.

10. Subramanian, I., Verma, S., Kumar, S., Jere, A. and Anamika, K. (2020) Multi-omics Data Integration, Interpretation, and Its Application. Bioinform Biol Insights, 14, 1177932219899051.

11. Goh, W.W.B., Wang, W. and Wong, L. (2017) Why Batch Effects Matter in Omics Data, and How to Avoid Them. Trends Biotechnol, 35, 498–507.

12. Leek, J.T., Scharpf, R.B., Bravo, H.C., Simcha, D., Langmead, B., Johnson, W.E., Geman, D., Baggerly, K. and Irizarry, R.A. (2010) Tackling the widespread and critical impact of batch effects in high-throughput data. Nat Rev Genet, 11, 733–739.

13. Han, W. and Li, L. (2022) Evaluating and minimizing batch effects in metabolomics. Mass Spectrom Rev, 41, 421–442.

14. Lazar, C., Meganck, S., Taminau, J., Steenhoff, D., Coletta, A., Molter, C., Weiss-Solís, D.Y., Duque, R., Bersini, H. and Nowé, A. (2013) Batch effect removal methods for microarray gene expression data integration: a survey. Brief Bioinform, 14, 469–490.

15. Molania, R., Foroutan, M., Gagnon-Bartsch, J.A., Gandolfo, L.C., Jain, A., Sinha, A., Olshansky, G., Dobrovic, A., Papenfuss, A.T. and Speed, T.P. (2023) Removing unwanted variation from large-scale RNA sequencing data with PRPS. Nat Biotechnol, 41, 82–95.

16. Nygaard, V., Rødland, E.A. and Hovig, E. (2016) Methods that remove batch effects while retaining group differences may lead to exaggerated confidence in downstream analyses. Biostatistics, 17, 29–39.

17. Adamer, M.F., Brüningk, S.C., Tejada-Arranz, A., Estermann, F., Basler, M. and Borgwardt, K. (2022) reComBat: batch-effect removal in large-scale multi-source gene-expression data integration. Bioinformatics Advances, 2, vbac071.

18. Chazarra-Gil, R., van Dongen, S., Kiselev, V.Y. and Hemberg, M. (2021) Flexible comparison of batch correction methods for single-cell RNA-seq using BatchBench. Nucleic Acids Res, 49, e42.

19. Johnson, W.E., Li, C. and Rabinovic, A. (2007) Adjusting batch effects in microarray expression data using empirical Bayes methods. Biostatistics, 8, 118–127.

20. Rong, Z., Tan, Q., Cao, L., Zhang, L., Deng, K., Huang, Y., Zhu, Z.-J., Li, Z. and Li, K. (2020) NormAE: Deep Adversarial Learning Model to Remove Batch Effects in Liquid Chromatography Mass Spectrometry-Based Metabolomics Data. Anal Chem, 92, 5082–5090.

21. Tran, H.T.N., Ang, K.S., Chevrier, M., Zhang, X., Lee, N.Y.S., Goh, M. and Chen, J. (2020) A benchmark of batch-effect correction methods for single-cell RNA sequencing data. Genome Biology, 21, 12.

22. Lakkis, J., Wang, D., Zhang, Y., Hu, G., Wang, K., Pan, H., Ungar, L., Reilly, M.P., Li, X. and Li, M. (2021) A joint deep learning model enables simultaneous batch effect correction, denoising, and clustering in single-cell transcriptomics. Genome Res, 31, 1753–1766.

23. Li, X., Wang, K., Lyu, Y., Pan, H., Zhang, J., Stambolian, D., Susztak, K., Reilly, M.P., Hu, G. and Li, M. (2020) Deep learning enables accurate clustering with batch effect removal in single-cell RNA-seq analysis. Nat Commun, 11, 2338.

24. Wang, T., Johnson, T.S., Shao, W., Lu, Z., Helm, B.R., Zhang, J. and Huang, K. (2019) BERMUDA: a novel deep transfer learning method for single-cell RNA sequencing batch correction reveals hidden high-resolution cellular subtypes. Genome Biol, 20, 165.

25. Shrikumar, A., Greenside, P. and Kundaje, A. (2019) Learning Important Features Through Propagating Activation Differences. 10.48550/arXiv.1704.02685.

26. Lundberg, S.M. and Lee, S.-I. (2017) A Unified Approach to Interpreting Model Predictions. In Advances in Neural Information Processing Systems. Curran Associates, Inc., Vol. 30.

27. Ulfenborg, B. (2019) Vertical and horizontal integration of multi-omics data with miodin. BMC Bioinformatics, 20, 649.

28. Bodein, A., Scott-Boyer, M.-P., Perin, O., Lê Cao, K.-A. and Droit, A. (2022) timeOmics: an R package for longitudinal multi-omics data integration. Bioinformatics, 38, 577–579.

29. Rau, A., Marot, G. and Jaffrézic, F. (2014) Differential meta-analysis of RNA-seq data from multiple studies. BMC Bioinformatics, 15, 91.

30. Lê Cao, K.-A., Boitard, S. and Besse, P. (2011) Sparse PLS discriminant analysis: biologically relevant feature selection and graphical displays for multiclass problems. BMC Bioinformatics, 12, 253.

31. Rohart, F., Eslami, A., Matigian, N., Bougeard, S. and Lê Cao, K.-A. (2017) MINT: a multivariate integrative method to identify reproducible molecular signatures across independent experiments and platforms. BMC Bioinformatics, 18, 128.

32. Moreno, P., Fexova, S., George, N., Manning, J.R., Miao, Z., Mohammed, S., Muñoz-Pomer, A., Fullgrabe, A., Bi, Y., Bush, N., et al. (2022) Expression Atlas update: gene and protein expression in multiple species. Nucleic Acids Research, 50, D129–D140.

33. Qi, H., Jiang, Z., Zhang, K., Yang, S., He, F. and Zhang, Z. (2018) PlaD: A Transcriptomics Database for Plant Defense Responses to Pathogens, Providing New Insights into Plant Immune System. Genomics Proteomics Bioinformatics, 16, 283–293.

34. Guimaraes, P.M., Guimaraes, L.A., Morgante, C.V., Silva, O.B., Araujo, A.C.G., Martins, A.C.Q., Saraiva, M.A.P., Oliveira, T.N., Togawa, R.C., Leal-Bertioli, S.C.M., et al. (2015) Root Transcriptome Analysis of Wild Peanut Reveals Candidate Genes for Nematode Resistance. PLoS ONE, 10, e0140937.

35. Mota, A.P.Z., Vidigal, B., Danchin, E.G.J., Togawa, R.C., Leal-Bertioli, S.C.M., Bertioli, D.J., Araujo, A.C.G., Brasileiro, A.C.M. and Guimaraes, P.M. (2018) Comparative root transcriptome of wild Arachis reveals NBS-LRR genes related to nematode resistance. BMC Plant Biology, 18, 159.

36. Vinson, C.C., Mota, A.P.Z., Oliveira, T.N., Guimaraes, L.A., Leal-Bertioli, S.C.M., Williams, T.C.R., Nepomuceno, A.L., Saraiva, M.A.P., Araujo, A.C.G., Guimaraes, P.M., et al. (2018) Early responses to dehydration in contrasting wild Arachis species. PLoS One, 13, e0198191.

37. Mota, A.P.Z., Brasileiro, A.C.M., Vidigal, B., Oliveira, T.N., da Cunha Quintana Martins, A., Saraiva, M.A. de P., de Araújo, A.C.G., Togawa, R.C., Grossi-de-Sá, M.F. and Guimaraes, P.M. (2021) Defining the combined stress response in wild Arachis. Sci Rep, 11, 11097.

38. Kornbrot, D. (2005) Point Biserial Correlation. In Encyclopedia of Statistics in Behavioral Science. John Wiley & Sons, Ltd.

39. Kruskal, W.H. and Wallis, W.A. (1952) Use of Ranks in One-Criterion Variance Analysis. Journal of the American Statistical Association, 47, 583–621.

40. Dunn, O.J. (1964) Multiple Comparisons Using Rank Sums. Technometrics, 6, 241–252.

41. Baldi, P. (2012) Autoencoders, Unsupervised Learning, and Deep Architectures. In Proceedings of ICML Workshop on Unsupervised and Transfer Learning. JMLR Workshop and Conference Proceedings, pp. 37–49.

42. Rumelhart, D.E., Hinton, G.E. and Williams, R.J. (1986) Learning Internal Representations by Error Propagation. 10.7551/mitpress/5236.003.0012.

43. Kingma, D.P. and Welling, M. (2022) Auto-Encoding Variational Bayes. 10.48550/arXiv.1312.6114.

44. Rezende, D.J., Mohamed, S. and Wierstra, D. (2014) Stochastic Backpropagation and Approximate Inference in Deep Generative Models. 10.48550/arXiv.1401.4082.

45. Pedregosa, F., Varoquaux, G., Gramfort, A., Michel, V., Thirion, B., Grisel, O., Blondel, M., Prettenhofer, P., Weiss, R., Dubourg, V., et al. (2011) Scikit-learn: Machine Learning in Python. MACHINE LEARNING IN PYTHON.

46. Abadi, M., Barham, P., Chen, J., Chen, Z., Davis, A., Dean, J., Devin, M., Ghemawat, S., Irving, G., Isard, M., et al. (2016) TensorFlow: a system for large-scale machine learning. In Proceedings of the 12th USENIX conference on Operating Systems Design and Implementation, OSDI’16. USENIX Association, USA, pp. 265–283.

47. Chollet, F. and others (2018) Keras: The Python Deep Learning library. Astrophysics Source Code Library.

48. Virtanen, P., Gommers, R., Oliphant, T.E., Haberland, M., Reddy, T., Cournapeau, D., Burovski, E., Peterson, P., Weckesser, W., Bright, J., et al. (2020) SciPy 1.0: fundamental algorithms for scientific computing in Python. Nat Methods, 17, 261–272.

49. Arlot, S. and Celisse, A. (2010) A survey of cross-validation procedures for model selection. Statist. Surv., 4.

50. Efron, B. and Tibshirani, R. (1997) Improvements on Cross-Validation: The .632+ Bootstrap Method. Journal of the American Statistical Association, 92, 548–560.

51. James, G., Witten, D., Hastie, T. and Tibshirani, R. (2013) Statistical Learning. In James, G., Witten, D., Hastie, T., Tibshirani, R. (eds), An Introduction to Statistical Learning: with Applications in R, Springer Texts in Statistics. Springer, New York, NY, pp. 15–57.

52. Refaeilzadeh, P., Tang, L. and Liu, H. (2009) Cross-Validation. In Liu, L., Özsu, M.T. (eds), Encyclopedia of Database Systems. Springer US, Boston, MA, pp. 532–538.

53. Breiman, L. (2001) Random Forests. Machine Learning, 45, 5–32.

54. Lundberg, S.M., Erion, G., Chen, H., DeGrave, A., Prutkin, J.M., Nair, B., Katz, R., Himmelfarb, J., Bansal, N. and Lee, S.-I. (2020) From local explanations to global understanding with explainable AI for trees. Nat Mach Intell, 2, 56–67.

55. R Core Team (2021) R: A Language and Environment for Statistical Computing R Foundation for Statistical Computing, Vienna, Austria.

56. Love, M.I., Huber, W. and Anders, S. (2014) Moderated estimation of fold change and dispersion for RNA-seq data with DESeq2. Genome Biol, 15, 550.

57. Rohart, F., Gautier, B., Singh, A. and Cao, K.-A.L. (2017) mixOmics: An R package for ‘omics feature selection and multiple data integration. PLOS Computational Biology, 13, e1005752.

58. Harris, C.R., Millman, K.J., Walt, S.J. van der, Gommers, R., Virtanen, P., Cournapeau, D., Wieser, E., Taylor, J., Berg, S., Smith, N.J., et al. (2020) Array programming with NumPy. Nature, 585, 357–362.

59. Tian, T., Liu, Y., Yan, H., You, Q., Yi, X., Du, Z., Xu, W. and Su, Z. (2017) agriGO v2.0: a GO analysis toolkit for the agricultural community, 2017 update. Nucleic Acids Res, 45, W122–W129.

60. Rhee, S.Y., Beavis, W., Berardini, T.Z., Chen, G., Dixon, D., Doyle, A., Garcia-Hernandez, M., Huala, E., Lander, G., Montoya, M., et al. (2003) The Arabidopsis Information Resource (TAIR): a model organism database providing a centralized, curated gateway to Arabidopsis biology, research materials and community. Nucleic Acids Research, 31, 224–228.

61. Schwacke, R., Ponce-Soto, G.Y., Krause, K., Bolger, A.M., Arsova, B., Hallab, A., Gruden, K., Stitt, M., Bolger, M.E. and Usadel, B. (2019) MapMan4: A Refined Protein Classification and Annotation Framework Applicable to Multi-Omics Data Analysis. Mol Plant, 12, 879–892.

62. Dash, S., Cannon, E.K.S., Kalberer, S.R., Farmer, A.D. and Cannon, S.B. (2016) Chapter 8 - PeanutBase and Other Bioinformatic Resources for Peanut. In Stalker, H.T., F. Wilson, R. (eds), Peanuts. AOCS Press, pp. 241–252.

63. Aurenhammer, F. (1991) Voronoi diagrams—a survey of a fundamental geometric data structure. ACM Comput. Surv., 23, 345–405.

64. Preparata, F.P. and Shamos, M.I. (1985) Computational Geometry Springer, New York, NY.

65. Edelsbrunner, H. and Seidel, R. (1986) Voronoi diagrams and arrangements. Discrete Comput Geom, 1, 25–44.

66. Aurenhammer, F., Klein, R. and Lee, D. (2013) Voronoi Diagrams And Delaunay Triangulations World Scientific Publishing Company.

67. Kolde, R. (2019) pheatmap: Pretty Heatmaps.

68. Huynh-Thu, V.A., Irrthum, A., Wehenkel, L. and Geurts, P. (2010) Inferring Regulatory Networks from Expression Data Using Tree-Based Methods. PLOS ONE, 5, e12776.

69. Jin, J., Tian, F., Yang, D.-C., Meng, Y.-Q., Kong, L., Luo, J. and Gao, G. (2017) PlantTFDB 4.0: toward a central hub for transcription factors and regulatory interactions in plants. Nucleic Acids Res, 45, D1040–D1045.

70. Tian, F., Yang, D.-C., Meng, Y.-Q., Jin, J. and Gao, G. (2020) PlantRegMap: charting functional regulatory maps in plants. Nucleic Acids Res, 48, D1104–D1113.

71. Hagberg, A.A., Schult, D.A. and Swart, P.J. (2008) Exploring Network Structure, Dynamics, and Function using NetworkX.

72. Guimaraes, P.M., Quintana, A.C., Mota, A.P.Z., Berbert, P.S., Ferreira, D. da S., de Aguiar, M.N., Pereira, B.M., de Araújo, A.C.G. and Brasileiro, A.C.M. (2022) Engineering Resistance against Sclerotinia sclerotiorum Using a Truncated NLR (TNx) and a Defense-Priming Gene. Plants (Basel), 11, 3483.

73. Mota, A.P.Z., Oliveira, T.N., Vinson, C.C., Williams, T.C.R., Costa, M.M. do C., Araujo, A.C.G., Danchin, E.G.J., Grossi-de-Sá, M.F., Guimaraes, P.M. and Brasileiro, A.C.M. (2019) Contrasting Effects of Wild Arachis Dehydrin Under Abiotic and Biotic Stresses. Front. Plant Sci., 10.

74. Clough, S.J. and Bent, A.F. (1998) Floral dip: a simplified method for Agrobacterium-mediated transformation of Arabidopsis thaliana. Plant J, 16, 735–743.

75. Vinson, C.C., Mota, A.P.Z., Porto, B.N., Oliveira, T.N., Sampaio, I., Lacerda, A.L., Danchin, E.G.J., Guimaraes, P.M., Williams, T.C.R. and Brasileiro, A.C.M. (2020) Characterization of raffinose metabolism genes uncovers a wild Arachis galactinol synthase conferring tolerance to abiotic stresses. Sci Rep, 10, 15258.

76. Júnior, J.D.A. de S., Coelho, R.R., Lourenço, I.T., Fragoso, R. da R., Viana, A.A.B., Macedo, L.L.P. de, Silva, M.C.M. da, Carneiro, R.M.G., Engler, G., Almeida-Engler, J. de, et al. (2013) Knocking-Down Meloidogyne incognita Proteases by Plant-Delivered dsRNA Has Negative Pleiotropic Effect on Nematode Vigor. PLOS ONE, 8, e85364.

77. Pratella, D., Ait-El-Mkadem Saadi, S., Bannwarth, S., Paquis-Fluckinger, V. and Bottini, S. (2021) A Survey of Autoencoder Algorithms to Pave the Diagnosis of Rare Diseases. Int J Mol Sci, 22, 10891.

78. Love, M.I., Huber, W. and Anders, S. (2014) Moderated estimation of fold change and dispersion for RNA-seq data with DESeq2. Genome Biol, 15, 550.

79. Perazzolli, M., Moretto, M., Fontana, P., Ferrarini, A., Velasco, R., Moser, C., Delledonne, M. and Pertot, I. (2012) Downy mildew resistance induced by Trichoderma harzianum T39 in susceptible grapevines partially mimics transcriptional changes of resistant genotypes. BMC Genomics, 13, 660.

80. Rodriguez, V.M., Padilla, G., Malvar, R.A., Kallenbach, M., Santiago, R. and Butrón, A. (2018) Maize Stem Response to Long-Term Attack by Sesamia nonagrioides. Front. Plant Sci., 9.

81. Vishwanathan, K., Zienkiewicz, K., Liu, Y., Janz, D., Feussner, I., Polle, A. and Haney, C.H. (2020) Ectomycorrhizal fungi induce systemic resistance against insects on a nonmycorrhizal plant in a CERK1-dependent manner. New Phytologist, 228, 728– 740.

82. Huang, H., Liu, B., Liu, L. and Song, S. (2017) Jasmonate action in plant growth and development. J Exp Bot, 68, 1349–1359.

83. Per, T.S., Khan, M.I.R., Anjum, N.A., Masood, A., Hussain, S.J. and Khan, N.A. (2018) Jasmonates in plants under abiotic stresses: Crosstalk with other phytohormones matters. Environmental and Experimental Botany, 145, 104–120.

84. Wang, Y., Mostafa, S., Zeng, W. and Jin, B. (2021) Function and Mechanism of Jasmonic Acid in Plant Responses to Abiotic and Biotic Stresses. Int J Mol Sci, 22, 8568.

85. Jones, J.D.G. and Dangl, J.L. (2006) The plant immune system. Nature, 444, 323–329.

86. Soltabayeva, A., Dauletova, N., Serik, S., Sandybek, M., Omondi, J.O., Kurmanbayeva, A. and Srivastava, S. (2022) Receptor-like Kinases (LRR-RLKs) in Response of Plants to Biotic and Abiotic Stresses. Plants (Basel), 11, 2660.

87. Takahashi, F. and Shinozaki, K. (2019) Long-distance signaling in plant stress response. Current Opinion in Plant Biology, 47, 106–111.

88. Tsuda, K. and Katagiri, F. (2010) Comparing signaling mechanisms engaged in pattern-triggered and effector-triggered immunity. Curr Opin Plant Biol, 13, 459–465.

89. McHale, L., Tan, X., Koehl, P. and Michelmore, R.W. (2006) Plant NBS-LRR proteins: adaptable guards. Genome Biology, 7, 212.

90. Takken, F.L.W. and Goverse, A. (2012) How to build a pathogen detector: structural basis of NB-LRR function. Curr Opin Plant Biol, 15, 375–384.

91. von Dahlen, J.K., Schulz, K., Nicolai, J. and Rose, L.E. (2023) Global expression patterns of R-genes in tomato and potato. Frontiers in Plant Science, 14.

92. Baillo, E.H., Kimotho, R.N., Zhang, Z. and Xu, P. (2019) Transcription Factors Associated with Abiotic and Biotic Stress Tolerance and Their Potential for Crops Improvement. Genes (Basel), 10, 771.

93. Lu, Q., Liu, H., Hong, Y., Liang, X., Li, S., Liu, H., Li, H., Wang, R., Deng, Q., Jiang, H., et al. (2022) Genome-Wide Identification and Expression of FAR1 Gene Family Provide Insight Into Pod Development in Peanut (Arachis hypogaea). Front. Plant Sci., 13.

94. Wang, Y., Zhang, M., Du, P., Liu, H., Zhang, Z., Xu, J., Qin, L., Huang, B., Zheng, Z., Dong, W., et al. (2022) Transcriptome analysis of pod mutant reveals plant hormones are important regulators in controlling pod size in peanut (Arachis hypogaea L.). PeerJ, 10, e12965.

